# A strategy for ionic current analysis of nanopore protein readouts

**DOI:** 10.64898/2026.09.28.755023

**Authors:** Andrew J. Stein, Neda Ghohabi Esfahani, Stuart Akeson, Pooria Daneshvar Kakhaki, Daphne Kontogiorgos-Heintz, Jeff Nivala, Miten Jain

## Abstract

**Background:** Nanopore-based protein reading holds promise for single-molecule proteomics, but analyzing the resulting ionic current signals presents substantial computational challenges. Unlike nucleic acids, peptide-associated ionic signals exhibit complex patterns with subtle transitions between amino acids that can be obscured by noise. We developed an integrated computational strategy for processing nanopore protein ionic current signals from synthetic protein constructs containing all 20 amino acids in controlled sequence contexts. We started with denoising to remove artifacts, followed by change point detection using variable-number and fixed-number segmentation methods.

**Results:** We employed dynamic time warping barycenter averaging to generate amino acid-specific reference templates. In parallel, we performed classification of amino acid regions using bidirectional LSTM networks trained on segment-level statistical features. Accuracy for 20-way amino acid classification remained limited, while charge-based binary classification reached 93.3% accuracy. Pairwise classification revealed that certain amino acids, particularly negatively charged residues, generate distinctive signatures. We used a profile hidden Markov model to capture sequential structure of protein translocation.

**Conclusion:** This approach provides computational strategies for automated segmentation, classification, and modeling of nanopore peptide translocation ionic current data.

## Background

Proteins are essential biomolecules that play crucial roles in bodily functions. Post-translational modifications, alternative splicing, and sequence variations give rise to many distinct proteoforms. These proteoforms contribute to the proteome’s dynamic and highly diverse nature, making single-molecule-level analysis essential for identifying variants[1, 2]. Neither proteome complexity nor quantity can be inferred from genomic or transcriptomic data alone, necessitating direct protein-level analysis. Existing methods do not fully capture the full heterogeneity of the proteome, leaving a significant proportion unexplored. Single-molecule mi protein sequencing technologies[3], especially nanopore-based approaches[3–5], could offer a path forward by enabling direct readout of individual intact protein molecules.

Nanopores have been employed for single-molecule DNA and RNA sequencing. In this process, molecules are threaded through the nanopore in a stepwise manner with the assistance of an enzymatic motor[6]. As the molecule translocates through the pore, it disrupts the baseline ionic current in a manner unique to the molecule at each step. These characteristic disruptions generate a signal that can be decoded to reveal the molecule’s sequence[7–9]. Translating this concept to polypeptides presents several fundamental challenges. Unlike nucleic acids with four bases, peptide analysis requires discriminating among 20 amino acids, each with distinct physicochemical properties[10]. Further, proteins are generally folded, contain heterogeneous non-uniform charges, complex three-dimensional structures, and post-translational modifications[11–14].

Despite these challenges, nanopores have shown sensitivity to amino acid regions in peptides, enabling applications like full-length protein reading[15, 16]. In nanopore-based protein reading, the generated fingerprint signal is used to identify proteins and detect deviations from expected signals[11]. However, achieving *de novo* protein sequencing and reconstructing a protein’s complete amino acid sequence remains challenging.

A fundamental challenge in nanopore protein analysis is accurate event detection and ionic current modeling of the resulting data[17]. This is achieved through segmentation[18], the process of identifying boundaries between step-wise ionic current steps, and machine learning (e.g., using HMMs, deep learning) to model ionic current segments[19–21]. For nucleic acids, segmentation methods have matured considerably, with approaches ranging from filter-derivative methods that detect abrupt signal changes, Bayesian methods that score boundaries using likelihood functions, to the most up-to-date Oxford Nanopore Dorado move-table[22–24]. Similarly, modeling can be performed using HMMs and transformers. However, these methods are not designed for peptide data that contain complex ionic current patterns with variable segment lengths.

Here, we present a computational strategy for nanopore protein ionic current signal analysis. This strategy includes strategies for denoising, segmentation, template construction, classification, and modeling of ionic current data. We developed a multi-phase denoising pipeline followed by change point detection using PELT and Dynamic Programming algorithms with custom cost functions tailored to protein signal characteristics. Following segmentation, we extracted statistical features from each segment and applied bidirectional LSTM networks for amino acid classification, achieving strong discrimination for several amino acid classes and revealing property-based groupings. We implemented DTW-based barycenter averaging to construct amino acid-specific templates and evaluated their discriminative capacity. Additionally, we explored HMM-based modeling to capture sequential dependencies in protein translocation. This integrated ionic current signal processing approach provides a foundation for analysis of nanopore protein sequencing data.

## Methods

### Nanopore Peptide Data Overview

Motone et al. developed a strategy for single-molecule analysis of long peptide strands[25] using a commercial Oxford Nanopore device (**Figure 1**). Briefly, peptide strands are loaded onto the nanopore array (**Figure 1.A**) and threaded through the nanopore using electrophoretic force (**Figure 1.B**). These peptide constructs contain a blocking domain that prevents full traversal through the pore. They are then enzymatically ratcheted using the translocase and unfoldase activity of the ClpX motor. This ClpX-mediated translocation is slower and produces reproducible ionic current signals (**Figure 1.C**). They made distinct protein constructs known as PASTORS (Proteins for Amino Acid Sequencing Through Optimized Regions) to demonstrate amino acid-level resolution in the ionic current data. Each construct was composed of five repeating sequence blocks (variable regions), each 59 amino acids long, based on a core sequence of glycine, serine, aspartic acid, and glutamic acid (**Figure 1.D**). A single, unique amino acid was introduced at the central position of each block, and the blocks were separated by two tyrosines (YY) to provide distinct ionic current bookends for downstream analysis. This strategic design enabled the analysis of five unique amino acids in a single protein strand. The eight PASTOR variants were synthesized to include all 20 amino acids appearing in two sequence contexts across the constructs. The PASTORs were named based on their amino acid combinations. This approach allowed for the investigation of amino acid properties in a controlled and systematic manner. Analysis of the PASTORs produced ionic current traces characterized by repetitive patterns. These patterns are attributed to the seven YY regions, visible as repeated dips in the signal (**Figure 1.D**). Between these dips, five distinctive and reproducible variations in the ionic current signals are observed, which correspond to the unique amino acid substitution within each variable region (VR). This opened new avenues to investigate these variable regions to analyze the ionic signatures associated with individual amino acid variations.

**Figure 1.**
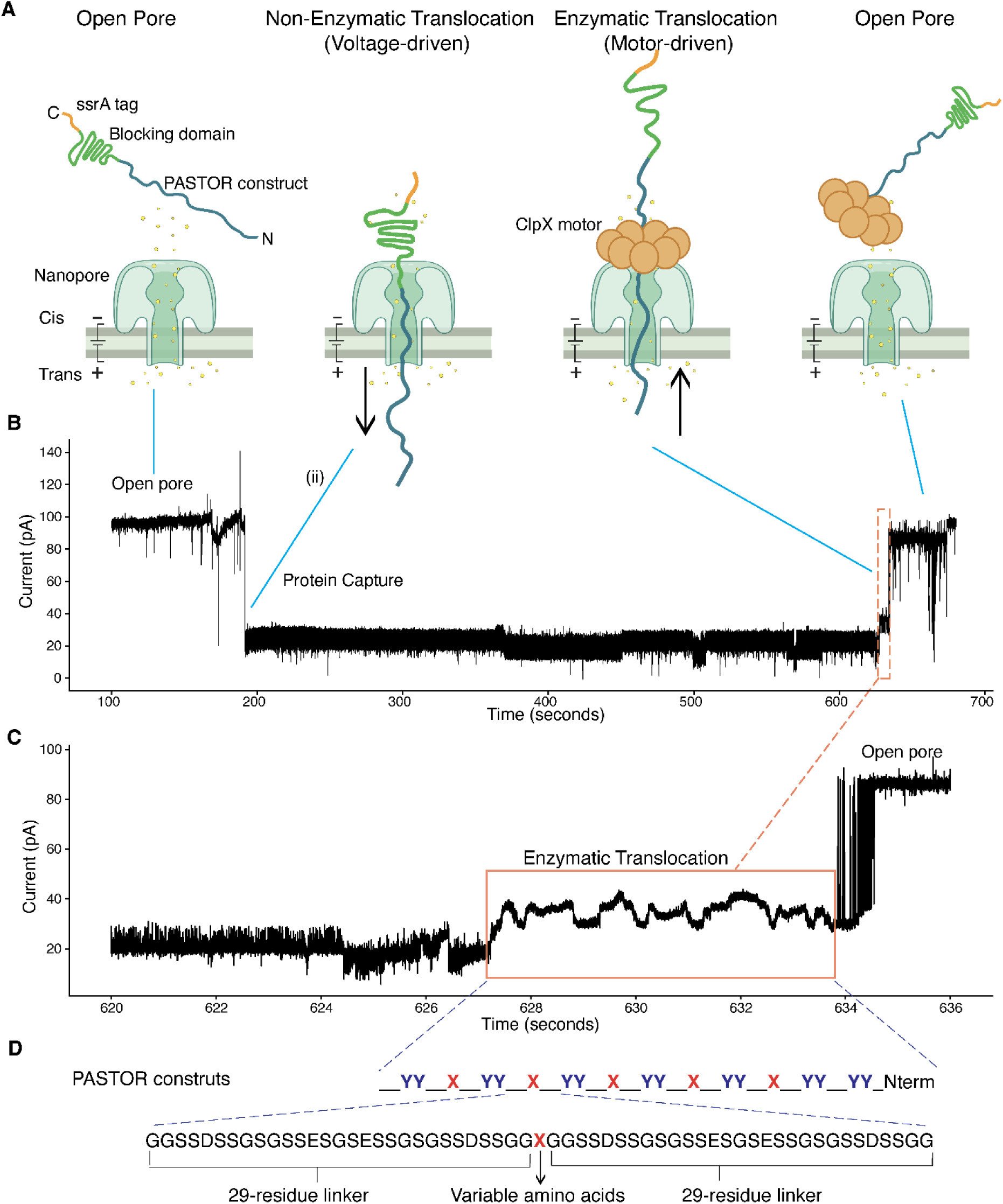
Nanopore-based protein reading using ClpX motor-assisted translocation and PASTOR constructs. **A.** Schematic of the nanopore protein sequencing workflow. Constructs contain an N-terminal ssrA tag for ClpX motor recognition, a blocking domain, and the PASTOR sequence. Left: Open pore state with high baseline ionic current. Second panel: Non-enzymatic voltage-driven translocation where the protein threads through the nanopore rapidly. Third panel: Enzymatic motor-driven translocation where the ClpX motor (orange hexamer) captures and unfolds the protein, then ratchets it through the nanopore in controlled steps, producing slower and more reproducible ionic current. Right: Return to open pore state after complete translocation. Cis and Trans indicate the two chambers separated by the membrane containing the nanopore. **B.** Representative ionic current trace showing the complete translocation event. The trace begins at open-pore baseline, drops sharply upon protein capture, maintains a reduced current level during enzymatic translocation, then returns to baseline. Time measured in seconds, current in picoamperes (pA). **C.** Magnified view of the enzymatic translocation phase (orange box from panel B) showing characteristic signal fluctuations as the protein passes through the nanopore. **D.** Structure of PASTOR constructs. Each PASTOR contains five variable regions separated by tyrosine-tyrosine (YY) markers (the 5 dip regions in signal). Variable regions are 59 amino acids long, built on a core sequence of glycine, serine, aspartic acid, and glutamic acid (G, S, D, E). A single unique amino acid (X, shown in red) is positioned at the center of each variable region. The construct has 29-residue linker sequences on both sides.

### Overview of the computational strategy

We developed an integrated computational strategy for processing nanopore protein ionic current signals from these synthetic polypeptide constructs containing all 20 amino acids in controlled sequence contexts (**Figure 2**).

**Figure 2.**
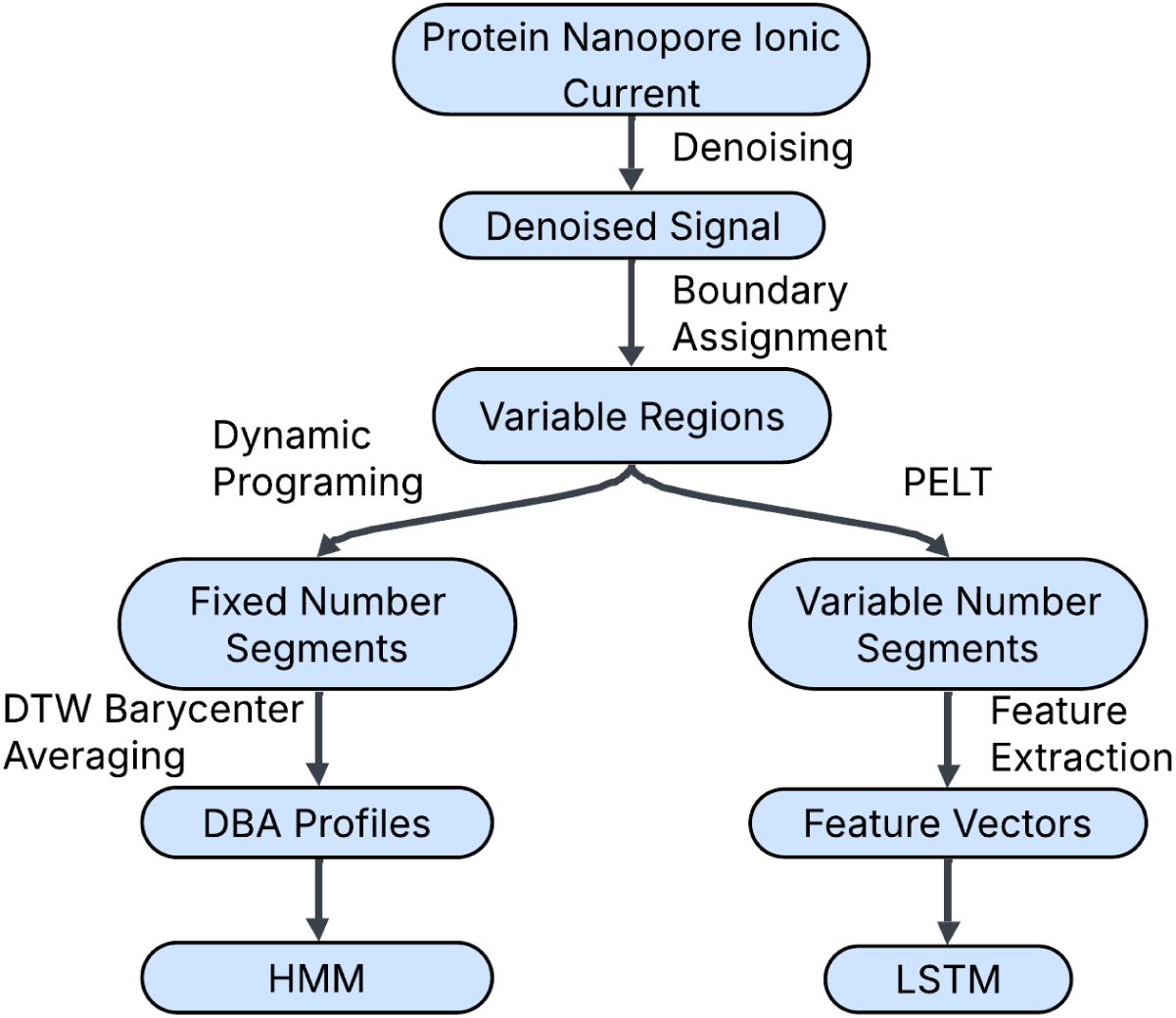
Overview of the computational strategy for processing nanopore protein ionic current signals.

### Data Preprocessing, Denoising and Boundary assignment

Raw ionic current signals were extracted from FAST5 files, and each PASTOR trace was denoised in three sequential passes, with each pass reconstructing the signal without the flagged samples using Huber regression.The first pass utilized an Isolation Forest algorithm to identify point anomalies, replacing samples whose anomaly score exceeded the contamination threshold. The second pass applied total variation (TV) denoising to suppress high-frequency noise; its L1 norm penalty on the signal gradient promotes finding piecewise-constant solutions, preserving the sharp transitions that downstream segmentation depends on.. The final pass utilizes continuous wavelet transform (CWT) analysis to flag residual current “spikes” in the signal, defined as exceeding the median by 0.25 times the Median Absolute Deviation (MAD). A single-sample dilation in both directions expands each identified point before reconstruction. Parameters for all three passes can be found in **Supplementary Table S1.** Variable region boundaries were determined by locating tyrosine-tyrosine (YY) markers, which produce characteristic signal dips in the ionic current trace. The absolute minimum within each of the YY dips was determined and assigned as the new boundary point between the adjacent variable regions.

### Segmentation strategy

We utilized offline change point detection algorithms to identify boundaries in ionic current signals corresponding to transitions between amino acid translocation events. These algorithms partition time-series data into segments with distinct statistical properties by identifying points where the signal characteristics shift. Change point detection methods consist of two key components: a search method that determines how candidate change points are evaluated and selected, and a cost function that quantifies the quality of a proposed segmentation. We evaluated multiple segmentation approaches, including Binary Segmentation, Window-Based and Bottom-Up Segmentation and focused on two methods that yielded reproducible segment counts across traces: Pruned Exact Linear Time (PELT) and Dynamic Programming (Dynp).

#### Pruned Exact Linear Time (PELT) Segmentation

PELT is an optimization-based algorithm that identifies the optimal set of change points by minimizing a cost function across the entire signal. The algorithm evaluates potential segmentation configurations and prunes suboptimal solutions early in the search process, making it computationally efficient for long signals. PELT determines the number of change points automatically based on a penalty parameter that controls the balance between the number of detected change points and segmentation quality. The algorithm operates by solving an optimization problem that minimizes the sum of segment costs plus a penalty term for the number of change points. Mathematically, this seeks to minimize [Σ *C*(*y*_{*t*_{*k* − 1}: *t*_*k*}) + β*K*] over all possible numbers of change points K and their positions {t_k}, where the sum runs from k=1 to K+1. Here, *C*(*y*_{*t*_{*k* − 1}: *t*_*k*}) represents the cost for the segment between consecutive change points t_{k-1} and t_k, β is the penalty parameter, and K is the total number of change points. The penalty term β directly controls segmentation sensitivity: higher values yield fewer, longer segments by penalizing each additional change point, while lower values permit finer segmentation. For our analysis, we used a penalty value of 5 and minimum segment size of 30 data points.

#### Dynamic Programming (Dynp) Segmentation

Dynamic Programming provides an alternative approach that requires specifying the exact number of change points. Dynp finds the optimal placement of a fixed number of breakpoints by exhaustively evaluating segmentation configurations. The algorithm minimizes the total cost across all segments. Specifically, it minimizes the sum Σ *C*(*y*_{*t*_{*k* − 1}: *t*_*k*}) over all change point positions {t_k}, where the sum runs from k=1 to n+1, and n is the specified number of breakpoints. This method was used when we required consistent segment counts across signals for comparative analysis.

#### Cost Function Design

The performance of change point detection depends on the cost function, which quantifies the homogeneity within segments and differences between them. We developed a variance-based cost function tailored to our protein ionic current data. This cost function calculates the variance within each segment as a measure of signal heterogeneity (Supplementary Algorithm 1). It centers the signal by subtracting its mean, then computes cumulative sums of the centered signal and its squared values to enable segment cost calculation. The scale parameter provides a multiplier for the variance, allowing adjustment of the cost magnitude relative to the penalty term. The minimum segment size (m=30) prevents oversegmentation by disallowing too short segments. The cost function preferentially places change points where signal variance shifts.

### Dynamic Time Warping and Barycenter Averaging

Following segmentation, all traces were standardized to produce exactly 35 segments using Dynamic Programming with a fixed number of breakpoints (n_bkps = 34). This uniform segment count was necessary for constructing representative amino acid templates via barycenter averaging. For each amino acid class, the segmented traces were normalized using z-score normalization. Each segment was then interpolated to a fixed length of 80 data points using linear interpolation, ensuring identical segment dimensionality for alignment.

#### DTW Barycenter Averaging (DBA)

To construct representative templates for each amino acid class, we employed DTW Barycenter Averaging (DBA). DBA computes a centroid sequence that minimizes the sum of squared DTW distances to all other data in that set. Unlike simple averaging, DBA iteratively refines an initial estimate by aligning all data to the current centroid estimate, computing aligned means, and updating the centroid. For each amino acid, all traces were stacked into a three-dimensional array. The DBA algorithm computed a barycenter of the same dimensionality (35 segments × 80 points), producing a single representative trace to capture the typical ionic current pattern for that amino acid across all observed translocation events.

#### Dynamic Time Warping Distance Metric

Dynamic Time Warping quantifies the similarity between two time series by finding the optimal alignment that minimizes the cumulative distance between matched points. DTW permits flexible temporal alignment, allowing segments with similar shapes but different speeds or phases to be recognized as similar. This is particularly relevant for nanopore signals where the same amino acid may translocate at slightly different rates. For two segments A and B of lengths m and n, DTW constructs a cost matrix C where each element C(i,j) represents the distance between points A(i) and B(j). The algorithm computes an accumulated cost matrix by dynamic programming, where each cell accumulates the minimum cost from three possible predecessor cells. The optimal alignment path through this matrix, backtracked from position (m,n) to (1,1), defines which points in A correspond to which points in B. The final DTW distance is the accumulated cost along this path, normalized by the path length to account for segment length differences. We implemented length-normalized DTW to enable fair comparisons across segments of varying duration. The raw DTW distance was divided by the average length of the two segments being compared: normalized_cost = DTW(A,B) / [(len(A) + len(B)) / 2].

#### Alignment Visualization and Quality Metrics

To evaluate alignment quality between the barycenter of a class and other traces of that class, we implemented a DTW visualization and quantification strategy. For each comparison, the fixed 35-segment barycenter and variable-length test trace were z-score normalized. A segment-level cost matrix was computed where element (i,j) represents the DTW distance between barycenter segment i and test segment j. The optimal alignment path was identified by dynamic programming, tracing back from the final accumulated cost to determine which barycenter segments aligned to which test segments. Three metrics quantified alignment quality. The total DTW cost is the sum of segment-level costs along the optimal alignment path, representing the cumulative dissimilarity between sequences. The mean cost per segment pair normalizes this total by dividing by the path length. The mean deviation from diagonal quantifies temporal distortion by averaging the absolute difference |i -j| across all path positions (i,j), where i is the barycenter segment index and j is the test segment index. Low deviation values indicate sequential alignment with minimal temporal warping, while high values indicate substantial reordering of segments was required to achieve optimal alignment.

#### Pairwise Inter/Intra Class Separation

To quantify how well each barycenter discriminates its own class from other classes, we computed pairwise separation ratios across all 20 classes. For each centroid-class pair, we calculated the median deviation from the diagonal when aligning that centroid to all traces of each class. The intra-class deviation represents the median alignment deviation between a centroid and traces from its own class, while the inter-class deviation represents the median alignment deviation between a centroid and traces from a different class. The separation ratio was defined as the inter-class deviation divided by the intra-class deviation. A separation ratio greater than 1.0 indicates that traces from the other class require more temporal warping to align with the centroid than traces from the centroid’s own class, suggesting distinguishable alignment signatures between the two classes. A ratio near or below 1.0 indicates that both classes align similarly to the centroid, implying poor discriminability. This analysis produced a 20×20 asymmetric matrix where entry (i,j) represents how well centroid i separates its own class from class j.

### Deep Learning Classification of Amino Acid Regions

Following variable-number segmentation, we extracted statistical features from each segment to characterize the ionic current distribution within that segment. For each segment, eight features were computed: skewness, kurtosis, standard deviation, maximum current, minimum current, median current, mean current, and segment length in data points. These features provided a numerical representation of each segment’s statistical properties.

#### LSTM Classifier Architecture

We employed a bidirectional Long Short-Term Memory (LSTM) network to classify amino acid regions based on their segmented ionic current traces. Each trace was represented as a sequence of feature vectors, where each vector contained the statistical features from one segment. The network architecture consisted of a two-layer bidirectional LSTM with 64 hidden units per direction, followed by a fully connected classification head. The bidirectional design processed sequences in both forward and reverse temporal directions, capturing dependencies between segments regardless of their position in the trace. LSTM layers included 30% dropout for regularization. The final hidden states from both directions were concatenated and passed through a two-layer fully connected network, 128 units with ReLU activation and 30% dropout, then an output layer with 20 units for amino acid classes. The model accepted variable-length sequences by using packed sequences to handle traces with variable numbers of segments.

#### Training Protocol and Model Evaluation

We employed stratified 5-fold cross-validation to evaluate classification performance. In each fold, the training set was used to compute class weights inversely proportional to class frequencies, which were applied in the cross-entropy loss function to prevent the model from being biased toward overrepresented amino acids. Models were trained for 100 epochs using the AdamW optimizer with learning rate 0.001 and weight decay 0.0001. Gradient clipping with maximum norm 1.0 prevented exploding gradients. Learning rate was reduced by 50% when validation loss plateaued for three consecutive evaluations. Sequences were processed in batches of 32, with padding applied to accommodate variable trace lengths within each batch. For each fold, we recorded validation set predictions and class probabilities. After completing all folds, predictions were aggregated to compute overall classification metrics including per-class accuracy, confusion matrix, classification report and multiclass ROC-AUC curves using one-vs-rest binarization.

#### Pairwise Classification

To assess discrimination capacity between specific amino acid pairs, we trained binary classifiers for all possible pairwise combinations of the 20 amino acids (190 total pairs). For each pair, we extracted traces belonging to only those two classes and relabeled them as binary targets. Each pairwise classifier used the same LSTM architecture but with a 2-unit output layer. Training followed an 80/20 train-test split with stratification to maintain class balance. Models were trained for 100 epochs with class-balanced cross-entropy loss. Pairwise accuracies were compiled into a symmetric 20×20 matrix.

### Hidden Markov Model

We built a profile hidden Markov model per class using the Yet Another hidden Markov model (YAHMM) package, which supports profile architectures combining both emitting and silent states, and allows the design of a linear topology. The model comprises four state types: **Match states** emit under a Gaussian whose mean is the DBA segment mean for that position and whose variance is the pileup variance across traces at that position. **Skip states** are silent and advance the path to a downstream match state without emissions, representing missing events in the query trace relative to the profile. **Slip states** are silent and return the path to an upstream match state, representing a repeated or missing event, expected when ClpX transiently releases the strand and it is re-threaded electrophoretically. **Interpolated states** (e.g. **Insert State**) may be inserted with emissions parameters averaged from the flanking profile states. Match states additionally permit self-loops. Traces were restricted to the interquartile range of trace length, then split 60/40 into profile constructions and inference sets. Each held-out trace was scored against every model by Viterbi decoding, with segment means scored against match-state Gaussians as univariate log-odds, and assigned to the highest-scoring model; classification was evaluated both pairwise and at the property-group level. Viterbi tracebacks were retained for inspection of candidate backslip events.

## Results

Accurate segmentation begins with effective signal preprocessing. Raw nanopore traces contain high-frequency noise that can mask genuine biological transitions. We developed a multi-phase denoising strategy that removes these artifacts while preserving the signal boundaries critical for detecting amino acid transitions. With denoised signals, segmentation algorithms can partition traces into regions corresponding to distinct translocation states. We implemented two complementary segmentation approaches. Variable-segment methods that adapt to natural heterogeneity in translocation dynamics, and fixed-segment methods that produce uniform outputs required for downstream modeling.

### Preprocessing, Denoising and Boundary Assignment of Raw Ionic Current Signals

Extraction of raw ionic current signals from FAST5 files yielded 670 PASTOR traces (see Methods), each containing five distinctive variable regions separated by tyrosine-tyrosine (YY) boundary markers (Figure 3A shows an example raw trace for PASTOR HDKER). We applied a multi-phase denoising strategy for cleaning the signal (see Methods). Variable region boundaries were assigned by identifying the midpoint of each YY dip (Figure 3B), ensuring complete capture of neighboring signal regions. This produced 3352 total variable regions with a median length of 2727 points. To ensure data quality, we excluded traces below the 5th percentile (1751 points) and above the 95th percentile (4570 points), keeping 2987 traces for subsequent segmentation.

**Figure 3.**
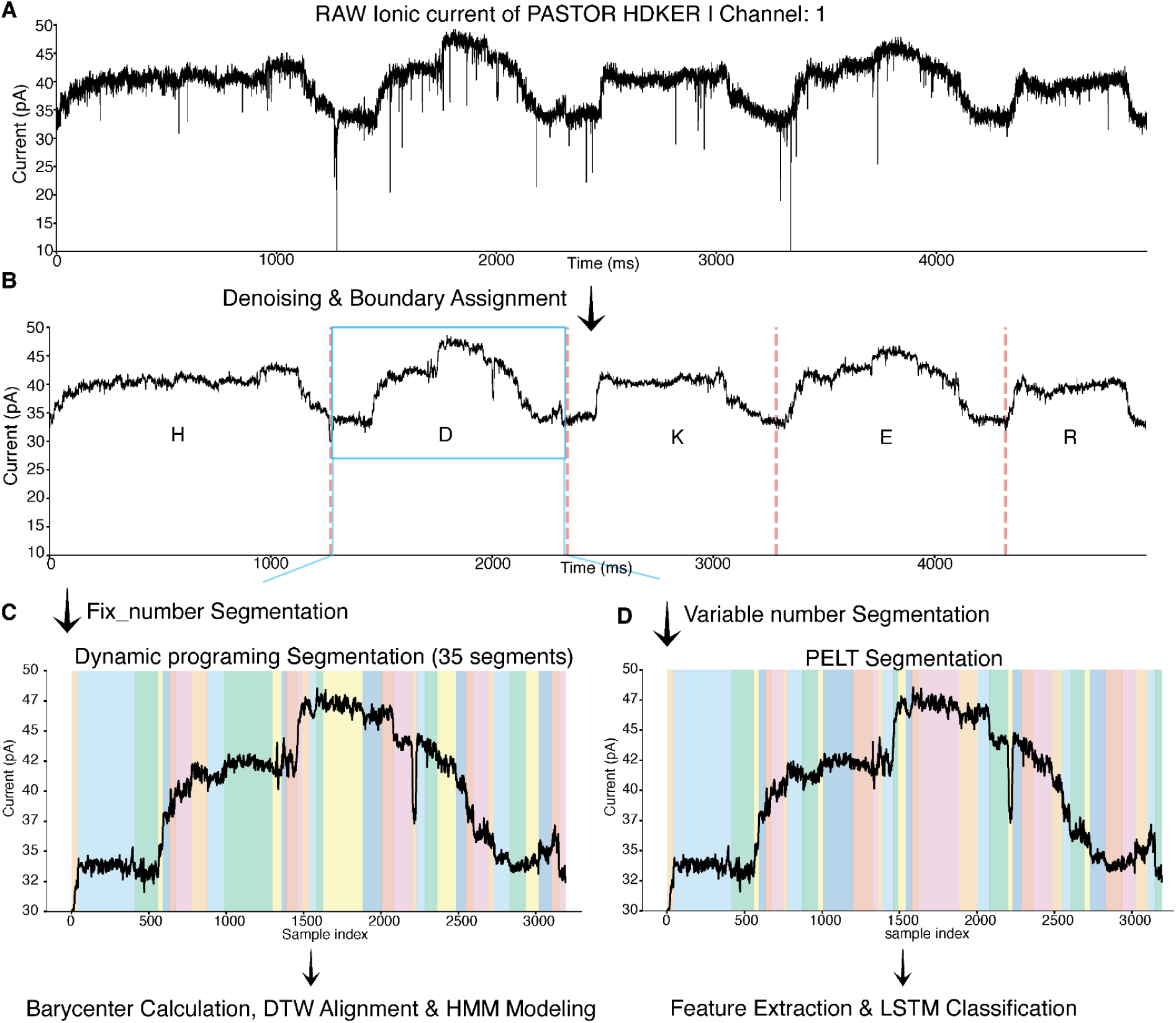
Ionic current signal processing strategy for nanopore protein readouts. **A.** Raw ionic current trace from PASTOR HDKER passing through the nanopore (Channel 1). The signal shows characteristic fluctuations as the protein translocates, with sharp dips corresponding to tyrosine-tyrosine boundary markers between variable regions. Current measured in picoamperes (pA) over time in milliseconds (ms). **B.** Denoised signal after applying the denoising strategy. Red dashed lines mark the boundaries of five variable regions, each labeled with its central amino acid (H, D, K, E, R). The blue box highlights the D region used in subsequent analysis steps. **C.** Fixed-segment approach using Dynamic Programming segmentation. The D region signal is divided into exactly 35 segments (shown in alternating colors). This produces consistent segment counts across all traces, which is required for barycenter construction. The X-axis shows sample index; Y-axis shows ionic current in pA. **D.** Variable-segment approach using PELT segmentation. The same D region is segmented based on detected change points in the signal, producing 38 segments. Colored bands indicate individual segments. X-axis shows sample index; the Y-axis shows ionic current in pA.

### Segmentation of Ionic Current Signals

Given the expectation for the polypeptide to traverse through the pore vestibule in a variable stepwise manner, we employed change point detection algorithms to identify boundaries in ionic current signals corresponding to transitions between amino acid translocation events. We applied two change point detection methods: PELT for variable-number segmentation and Dynamic Programming (Dynp) for fixed-number segmentation (see Methods).

### Variable-Number Segmentation

PELT segmentation was performed to identify boundaries between distinct current levels within each trace. An example segmentation for amino acid D is shown in **Figure 3D**.

PELT produced a median of 35 segments across all traces (**Supplementary Figure 1.A** shows the distribution of the number of segments across all variable regions). As expected from the inherent time variability in nanopore translocation, longer traces yielded more segments while shorter traces produced fewer (**Supplementary Figure 1.B** shows a 2D histogram of segment count vs. trace length; Spearman ρ = 0.51, p < 10⁻¹⁹⁵), demonstrating a moderate positive correlation between trace length and segmentation. While trace length explains approximately 30% of the variance in segment count (Pearson r² ∼0.31), the moderate correlation suggests that PELT segmentation may capture structural features in the ionic current signal beyond simply dividing longer traces into proportionally more pieces. This indicates that other factors, including signal quality, amino acid composition, and biological variability in translocation dynamics also contribute to determining segment boundaries.

The distribution of segment counts was consistent across the eight unique PASTOR constructs (**Supplementary Figure 1.C**), indicating consistency and reproducibility in segmentation. Visual inspection of traces from the same construct and across different constructs confirmed segmentation consistency (**Supplementary Figures 2 & 3** show segmentation of all five amino acid regions from the eight PASTORs, along with three example traces from within the same PASTOR class). Also, the median of 35 segments is aligned with the reported average of 2 amino acids per translocation step of the motor protein.

### Fixed-Number Segmentation

For downstream methods requiring fixed-length feature vectors (reference profiles), we implemented Dynamic Programming (Dynp) based segmentation using the median segment count (35) derived from PELT analysis. Using 34 breakpoints and the custom variance-based cost function (see Methods), Dynp produced exactly 35 segments for all traces (Figure 3C shows an example for amino acid D).

### DTW Barycenter Alignment Separation Performance of Amino Acid Regions

To create representative templates for each amino acid class, we constructed barycenters using DTW barycenter averaging on the Dynp segmented traces. All traces were z-score normalized to remove baseline differences while preserving within-segment current variations. Each segment was interpolated to 80 data points to ensure uniform dimensionality for alignment (see Methods). This procedure generated 20 barycenters, one for each amino acid (**Figure 4A** shows the barycenter for amino acid D; **Supplementary Figure 4** shows all 20 barycenters on top of each other).

**Figure 4.**
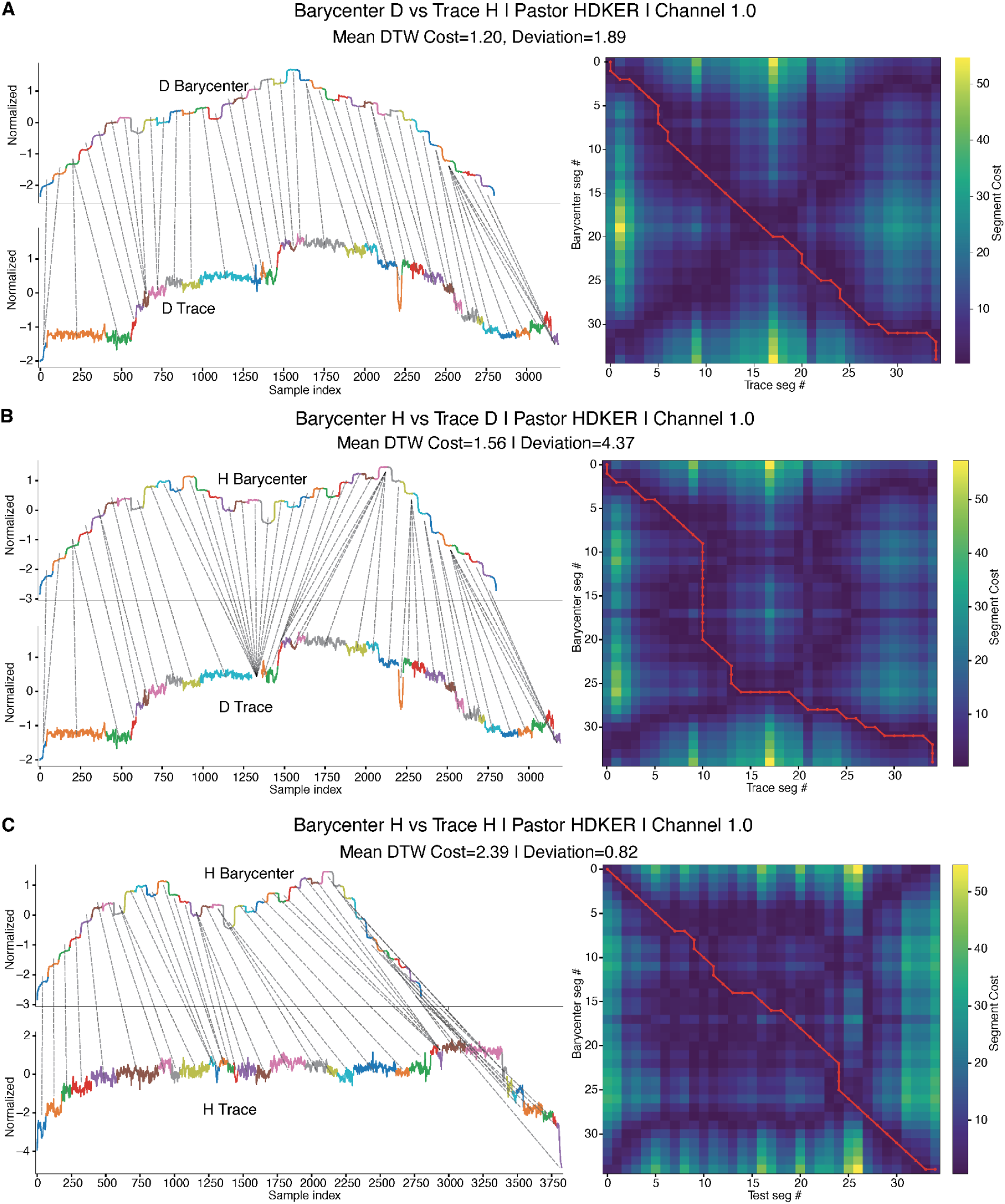
DTW barycenter alignment for intra-class and cross-class comparisons. **A.** DTW alignment between amino acid D barycenter (top) and an individual D trace from PASTOR HDKER, Channel 1 (bottom), both signals z-score normalized. The barycenter represents the average signal pattern for amino acid D calculated from multiple traces. Gray connection lines indicate which segments align between the two sequences based on the optimal DTW path. The cost matrix heatmap (right) displays the segment-wise DTW distance, with the red line showing the optimal alignment path. Mean DTW cost per segment = 1.20, Path deviation from diagonal = 1.89. **B.** DTW alignment between amino acid H barycenter (top) and the same individual D trace from panel A (bottom). The cost matrix shows the segment-wise DTW distance with the red line indicating the optimal alignment path. Mean DTW cost = 1.56, Path deviation = 4.37. DTW alignment between amino acid H barycenter (top) and an individual H trace from PASTOR HDKER, Channel 1 (bottom), both z-score normalized. The cost matrix shows segment-wise DTW distances with the optimal alignment path shown in red. Mean DTW cost = 2.39, Path deviation from diagonal = 0.82.

To assess how well barycenters distinguish amino acid classes, we computed pairwise separation ratios by comparing intra-class distances (barycenter to same-class traces) with inter-class distances (barycenter to different-class traces). Using path deviation as the metric, 90% of amino acid pairs showed separation ratios above 1, with a median of 1.27 (**Supplementary Figure 5A** shows the full separation matrix; **Supplementary Figure 5B** shows the distribution). The best-separated pairs exhibited ratios above 2, including D centroid versus C traces (2.22), D versus E traces (2.01), and several pairs involving L, P, V, and W centroids. These high ratios indicated that cross-class traces require substantially more temporal warping to align with the wrong barycenter compared to same-class alignments, demonstrating template discrimination. Conversely, poorly separated pairs showed ratios near or below 1.0, particularly involving similar amino acids. For example, E centroid versus D traces (0.79) and C centroid versus A traces (0.91) demonstrated overlap in alignment characteristics. The separation analysis reveals that while many amino acid pairs show clear barycenter-based discrimination, systematic overlap exists among physicochemically similar residues.

Visual inspection of DTW alignments provided insight into barycenter-trace correspondence. Intra-class alignments (barycenter and trace from the same amino acid class) generally showed alignment paths close to the diagonal with moderate costs. For example, alignment of D barycenter to a D trace yielded a mean DTW cost of 1.20 with a path deviation of 1.89 (**Figure 4A**), while H barycenter to H trace alignment produced a mean cost of 2.39 with a deviation of 0.82 (**Figure 4C**). These metrics varied across amino acid classes, reflecting differences within each class. Cross-class alignments (barycenter from one amino acid aligned to a trace from a different amino acid) showed distinct alignment characteristics. Aligning H barycenter to a D trace resulted in a mean cost of 1.56 but substantially higher path deviation of 4.37 (**Figure 4B**). While DTW’s flexible matching properties allow alignment between any two sequences, cross-class comparisons showed either higher costs, increased path deviations, or both compared to intra-class alignments. These suggest that the ionic current signal of different classes can differ enough to cause a higher cost when aligned to traces from other classes. However, the overlap in cost and deviation distributions between intra- and cross-class alignments shows that DTW metrics alone provide limited class discrimination for some amino acid pairs.

These observations motivated exploration of hidden Markov model approaches, which explicitly model sequential dependencies and class-specific emission properties rather than relying only on template distance metrics.

### LSTM-Based Amino Acid Region Classification

To evaluate the discriminative capacity of segment-level features, we trained a bidirectional LSTM classifier on statistical features extracted from PELT-segmented traces. Eight normalized features were computed per segment: skewness, kurtosis, standard deviation, maximum, minimum, median, mean, and segment length. These feature sequences were fed into a two-layer bidirectional LSTM network with 64 hidden units per direction, followed by a fully connected classification head (see Methods).

### 20-Way Multiclass Classification

Five-fold cross-validation on all 20 amino acids yielded an average accuracy of 20.7% ± 2.9%, in comparison to 5% random (**Supplementary Table S2** shows overall and per-class classification metrics including precision, recall, f1 score). Training dynamics showed consistent convergence across all folds, with validation loss plateauing after approximately 70 epochs (**Supplementary Figure 6** shows representative training curves). Validation accuracy increased progressively from ∼5% to ∼25% over 100 epochs, stabilizing in the later training phase. The adaptive learning rate scheduler reduced the learning rate from 10⁻³ to ∼10⁻⁶, enabling fine-tuned optimization. Training and validation curves tracked closely, indicating that performance was limited by the discriminative information rather than by overfitting.

Individual amino acid classes showed markedly different classification performance (**Supplementary Figure 7. A** shows ROC curves for all 20 classes; **Supplementary Figure 7. B** shows the confusion matrix). Amino acid D achieved the highest per-class accuracy at 81%, with an ROC AUC of 0.97, indicating highly distinctive ionic current signatures. Amino acids E (54.1% accuracy, AUC = 0.93) and R (47.2% accuracy, AUC = 0.88) also demonstrated strong classification performance. Amino acid A achieved 41.4% accuracy with an AUC of 0.85, showing good discriminability despite moderate recall. In contrast, several amino acids showed much lower individual accuracies, including C (1.5%), I (4.1%), V (5.8%) and N (6.0%). The confusion matrix revealed that misclassifications were not random but concentrated among amino acids with similar physicochemical properties or size characteristics. For instance, amino acids within the same size class or polarity group showed higher confusion rates, suggesting that these properties influence ionic current patterns in overlapping ways.

### Pairwise Classification

To further assess discrimination between specific amino acid pairs, we trained binary classifiers for all 190 possible pairwise combinations (**Figure 5** shows the pairwise accuracy heatmap). Pairwise accuracies ranged from 49.0% to 100%, with an average of approximately 75%, higher than the 20-way classification performance. This indicated that while distinguishing among all 20 amino acids simultaneously is challenging, many pairs are readily separable.

**Figure 5.**
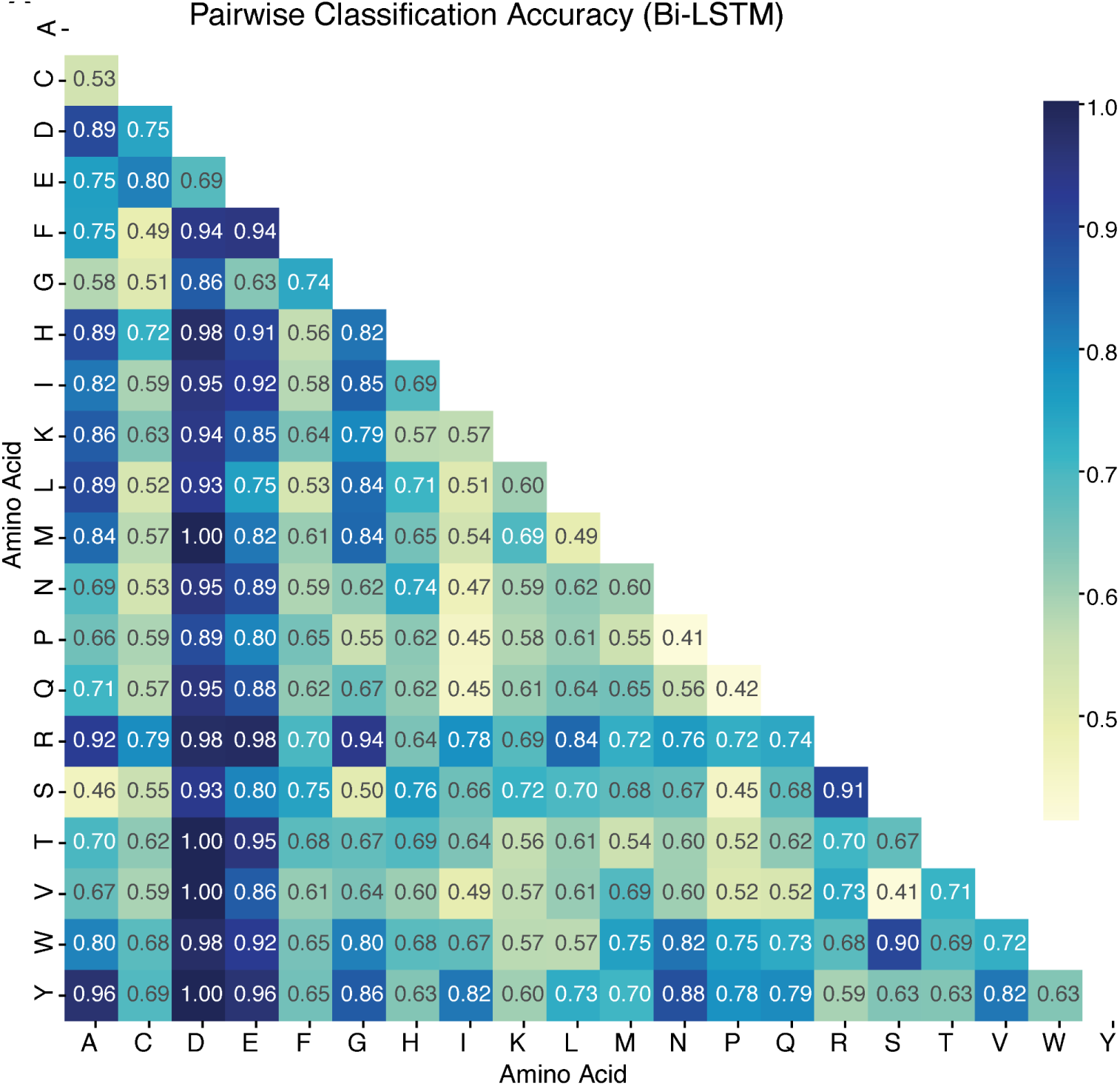
Pairwise LSTM classification. Eight statistical features (standard deviation, mean, median, maximum, minimum, duration, skewness, kurtosis) are calculated from each segment of PELT-segmented traces. These feature sequences are fed into a bidirectional LSTM network for classification. A. The heatmap shows pairwise classification accuracy between all 20 amino acids (190 total comparisons). Dark blue indicates high accuracy, while lighter blue shows lower accuracy.

Amino acid D showed particularly strong pairwise discrimination, achieving >93% accuracy against 15 of the 19 other amino acids, with perfect (100%) separation from M, T, V, and Y. Similarly, amino acid E demonstrated >85% accuracy against 14 other amino acids, with particularly high discrimination from charged and large amino acids (>94% vs. D, >98% vs. R). The charged amino acid R also showed robust pairwise performance, achieving >86% accuracy against 11 other amino acids. Several amino acid pairs showed near-perfect discrimination, including D vs. M (100%), D vs. T (100%), D vs. V (100%), D vs. Y (100%), and E vs. R (98.3%). These high accuracies suggest that certain amino acids generate highly distinctive translocation signatures that are consistently recognizable across different sequence contexts. Conversely, some pairs showed lower discrimination, particularly among amino acids with similar properties. For example, C vs. F (49.0%), C vs. L (52.5%), and F vs. L (52.5%) demonstrated near-chance performance, consistent with their shared hydrophobic character and similar molecular volumes. The pairwise heatmap revealed clustering of confusion among small nonpolar amino acids and among medium-sized amino acids, reflecting the challenge of discriminating between amino acids with overlapping biophysical properties from ionic current.

These results demonstrated that while certain amino acids produce highly characteristic signals enabling robust classification, improved discrimination of similar amino acids will likely require expanded training datasets, optimization of construct designs to enhance signal diversity, and potentially integration of additional signal features or contextual information from neighboring amino acids.

### Property-Based Multi-Group Classification

To assess whether amino acid groupings based on physicochemical properties enable more robust classification than individual amino acid identification, we trained LSTM classifiers on groups defined by charge, size, and polarity characteristics.

### Charge-Based Classification

Binary classification of positively charged (K, R, H; n=387) versus negatively charged (D, E; n=267) amino acids achieved the highest performance among all multi-group tasks, with an average accuracy of 93.3% ± 1.3% across five-fold cross-validation (**Supplementary Table S3** shows detailed metrics). The classifier demonstrated balanced performance between both classes, correctly identifying 93.0% of positive and 93.6% of negative amino acids. ROC analysis yielded an AUC of 0.963, indicating near-complete separation based on charge properties (**Supplementary Figure 8.A** shows the ROC curve and confusion matrix). This demonstrates that the presence of charge can be associated with distinctive ionic current signatures detectable by the model.

Expanding to three-way classification, positive (K, R, H), negative (D, E), and neutral (15 remaining amino acids; n=2,333), yielded 70.0% ± 1.9% accuracy (**Supplementary Table S4**). Negative amino acids maintained high recall (88%), while positive amino acids showed 65.6% accuracy with moderate precision (31%). The neutral class, despite comprising 78% of samples, achieved 68.7% accuracy but was frequently confused with charged amino acids (**Supplementary Figure 8.B** shows the confusion matrix). Multi-class ROC analysis produced macro-averaged AUC of 0.852, with negative amino acids showing the strongest discriminative capacity (AUC = 0.950) followed by neutral (AUC = 0.814) and positive (AUC = 0.793) groups. **Supplementary Figure8.B** shows one-vs-rest ROC curves.

### Size-Based Classification

Five-way classification based on molecular size, very small (G, A, S), small (C, D, N, P, T, V), medium (E, I, L, Q), large (H, K, M, F), and very large (R, W, Y), achieved 36.0% ± 1.6% accuracy (**Supplementary Table S5**). Performance varied substantially across size categories, with very small amino acids showing the highest accuracy at 75.7% and strong recall (76%). Very large amino acids achieved 56.4% accuracy with 56% recall. In contrast, medium-sized amino acids showed only 8.2% accuracy, with most misclassifications distributed among neighboring size categories (**Supplementary Figure 8.C** shows the confusion matrix). This pattern suggests that while extreme size differences produce distinguishable signals, intermediate sizes generate overlapping current distributions. Macro-averaged ROC AUC was 0.715, with very small (AUC = 0.831) and very large (AUC = 0.832) categories demonstrating the strongest discrimination. **Supplementary Figure 8.C** shows size-based ROC curves.

### Polarity-Based Classification

Four-way classification by polarity (nonpolar (A, G, I, L, M, F, W, P, V), polar (S, T, N, Q, C, Y), positive (K, R, H), and negative (D, E)) yielded 49.5% ± 1.5% accuracy (**Supplementary Table S6**). Negative amino acids again showed the highest per-class accuracy (88.0%) and strong precision-recall balance (F1 = 0.70). Nonpolar amino acids achieved 60.0% accuracy with balanced precision and recall (F1 = 0.60). However, polar amino acids showed substantially lower performance (16.4% accuracy, F1 = 0.23), with frequent misclassification as nonpolar, suggesting overlapping signal characteristics between these categories. Positive amino acids demonstrated 65.1% accuracy with higher recall (65%) than precision (32%), indicating a tendency for the model to over-predict this category. Macro-averaged ROC AUC was 0.747, with negative amino acids showing the strongest discrimination (AUC = 0.953). **Supplementary Figure 8. D** shows polarity-based ROC curves and confusion matrix.

These results indicate that amino acid charge produces the most discriminative ionic current signatures, followed by extreme size differences. The consistently strong performance of negatively charged amino acids (D, E) across all classification schemes aligns with their high individual classification accuracies observed in the 20-way task, suggesting these amino acids generate uniquely identifiable translocation patterns. The moderate performance for intermediate categories (medium-sized, polar) reflects the challenge of distinguishing amino acids with overlapping physicochemical properties, consistent with the pairwise classification results.

### Hidden Markov Modeling

We employed a profile hidden Markov model to capture the sequential structure of protein translocation under the assumption of a set step-size (**Supplementary Figure 9**). HMMs provide a natural framework for modeling the stepwise motor mechanism, explicitly representing state transitions and amino acid-specific emission properties. The ability to model state transitions allows the HMM to model motor or strand translocation mechanics, such as backslips from the motor or stalls in the pore. This probabilistic approach complements the pattern recognition strengths of deep learning with explicit modeling of biological constraints. To ensure robust modeling, we restricted our dataset to a clean subset of traces falling within the interquartile range (IQR) of all trace lengths. Classification was performed both at the pairwise amino acid level (**Supplementary Figure 10**) and at the domain level to enable direct comparison with the LSTM.

At the coarsest level, binary discrimination between positively and negatively charged residues achieved 83.9% accuracy (**Figure 6A**, **left**). Visual examination of the Viterbi path tracebacks revealed clear separation between categories: a given amino acid scored substantially higher against its own model (**H**, log probability = −199.1) than against a model from the opposing charge class (**D**, log probability = −347.4) (**Figure 6A**, **right**). Expanding to three-way classification by charge (positive, negative, and neutral), accuracy remained strong at 72.7% (**Figure 6B**, **left**). Path tracebacks showed that comparing a neutral amino acid to a negatively charged model produced a large deviation (**D**, log probability = −73.3), whereas comparison to a positively charged model yielded less separation (**H and V**, log probabilities = −34.2 and −25.8, respectively).

**Figure 6.**
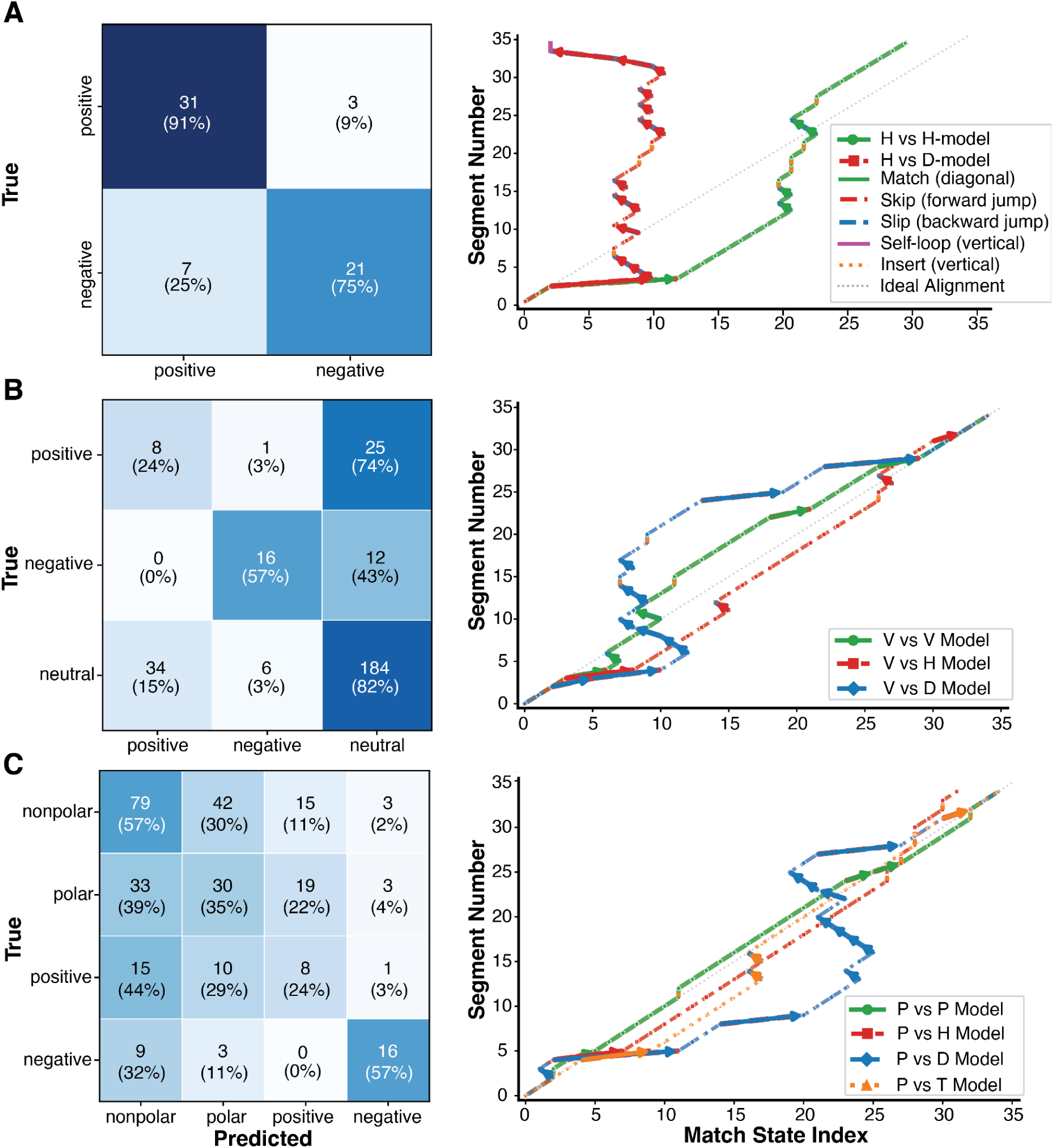
Hidden Markov model inference. **A. Left:** Binary charge-based classification (Positive vs. Negative). Confusion matrix showing classification of positively charged amino acids (K, R, H; n=28) versus negatively charged amino acids (D, E; n=34). Numbers indicate the count of samples and predictions in each category, with darker colors indicating higher counts. **Right:** Model traversal of amino acid H (positive) vs. its own model (green), and H vs. amino acid D (negative, red). The x-axis is the match state index in the profile, the y-axis is the segment index. A diagonal move denotes the model matching at a given state and progressing forward to the next. A move to the left denotes a “backslip” where the model moved to a previous match state to match a segment. A move right is the opposite, a “skip” where the model skipped some match states to match a downstream state. **B. Left:** Three-way charge-based classification (Positive vs. Negative vs. Neutral). Confusion matrix for classification of positively charged (K, R, H; n=28), negatively charged (D, E; n=34), and neutral amino acids (15 remaining amino acids; n=224). **Right:** Model traversal of amino acid V (neutral) vs. its own model (green), and V vs. H (positive, red), and V vs. D (negative, blue). **C. Left:** Four-way polarity-based classification. Confusion matrix for classification by polarity: Nonpolar (A, G, I, L, M, F, W, P, V; n=139), Polar (S, T, N, Q, C, Y; n=85), Positive (K, R, H; n=28), and Negative (D, E; n=34). **Right:** Model traversal of amino acid P (nonpolar) vs. its own model (green), and P vs. H (positive, red), P vs. D (negative, blue), and P vs. T (polar, orange).

When classifying by polarity, accuracy was 46.5% (**Figure 6C**, **left**). The nonpolar amino acid proline exhibited a distinct traceback relative to negatively charged models (**D**, log probability = −75.7) but yielded nearly identical log probabilities when compared to its own class (**P**, log probability = −29.5) and to a polar residue (**T**, log probability = −29.9), illustrating the source of confusion between nonpolar and polar categories.

We noticed that classifying based on size decreased accuracy to 35.0% (**Supplementary Figure 11**, **left**), reflecting the increased difficulty of distinguishing residues that differ primarily in size rather than in charge or polarity. Traceback analysis corroborated this difficulty: a very large amino acid (**W**, log probability = −37.4) produced a traceback similar to that of the large class (**H**, log probability = −57.6) but diverged more clearly from the medium (**E**, log probability = −83.0), small (**D**, log probability = −79.2), and very small (**S**, log probability = −82.6) classes (**Supplementary Figure 11, right**).

These results demonstrate that while the profile HMM sacrifices some accuracy relative to the nonlinear LSTM, it achieves comparable performance at the domain level while providing an interpretable, generative model of the translocation process.

### Profiling Putative Backslips with profile HMMs

Beyond classification, the profile HMM provides an interpretable linear model whose Viterbi path tracebacks can be visualized directly (as shown above) or leveraged for further analysis. We applied the model to profile potential backslip events during translocation. The model’s state architecture (**Supplementary Figure 9**) accommodates both skip transitions, analogous to deletion events in a conventional multiple-sequence alignment (MSA) HMM, and backslip transitions, analogous to insertions. Additionally, the model permits self-loops on match states and can insert interpolated states whose emission parameters are derived from the flanking profile states. This design was motivated by the need to capture pore-level events such as the transient loss of a peptide by the ClpX enzyme, followed by electrophoretic re-threading of the strand. Examples of these behaviors are shown in **Figure 7**. Between profile states 26 and 27 (**Figure 7A**), the traceback reveals a characteristic backslip signature: the decoded path advances to a downstream state before returning to an upstream one, producing a visible reversal in ionic current space. A related pattern is evident in **Figure 7B**, where profile state 11 was initially skipped but then visited following a backslip after state 13. **Supplementary Figure 12** presents a trace in which repeated backslips occur early in the state sequence and persist throughout, manifesting as repeated blocks of similar ionic current levels, but demonstrating that these models are still susceptible to being biased by noisy traces.

**Figure 7.**
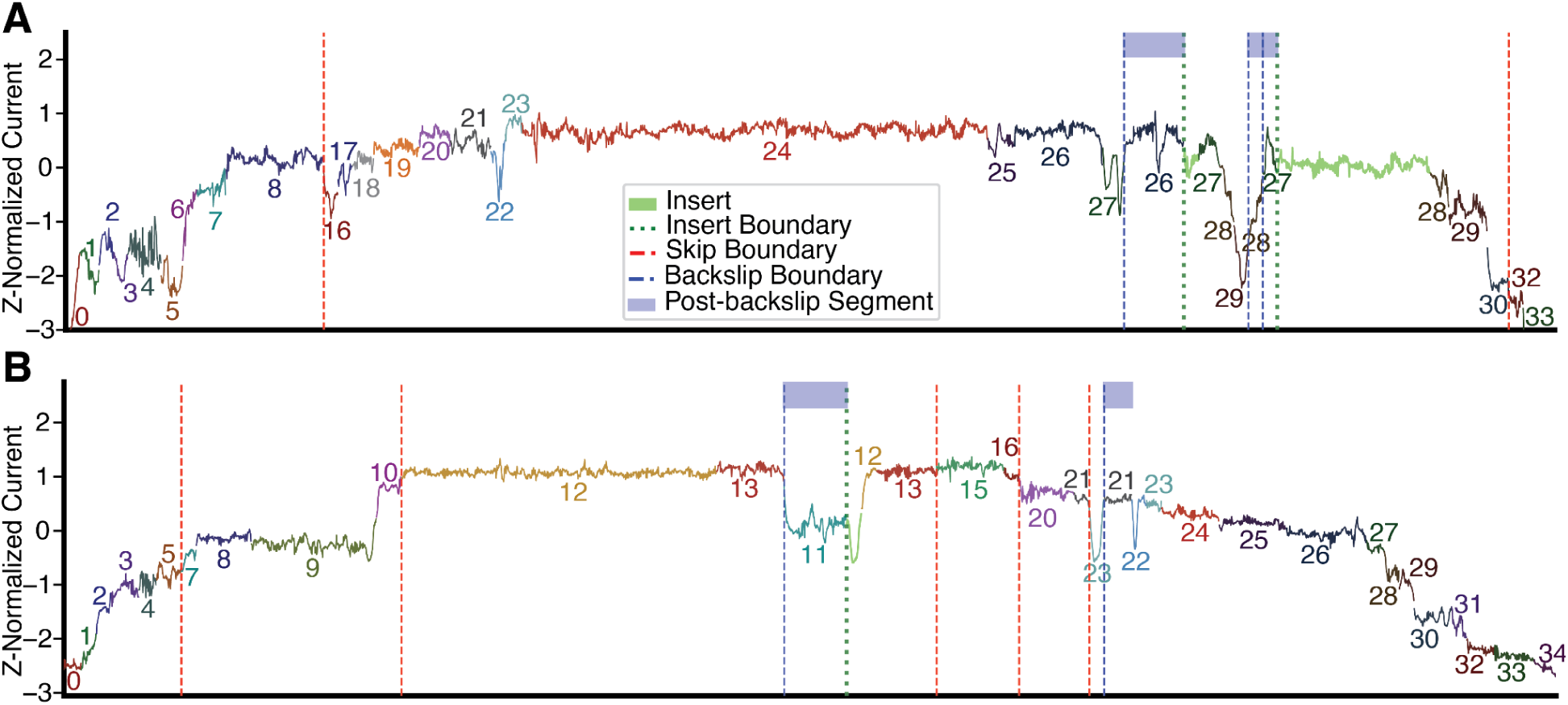
HMM Inference on Segmented Ionic Current Traces. **A.** Segmented Ionic current trace (G) colored to represent HMM Inference. The count and color sequence represents each unique match state the model traversed when inferring the trace. The red dashed line represents a point where the model chose to skip model match states until a downstream match state. The blue dashed line represents a point where the model slipped backwards to an upstream match state. The green dashed line and following segment represents the model inserting a state with a mean and variance averaged between the preceding and following states. The y-axis is z-score-normalized ionic current in picoamperes. **B.** Segmented Ionic current trace (D) colored to represent HMM inference.

In the absence of ground-truth backslip annotations, we cannot validate these assignments directly. Nevertheless, the specialized architecture of this profile HMM enables identification of candidate backslip events in ionic current data, highlighting regions of interest for further experimental investigation.

## Discussion

We developed a computational strategy for analyzing nanopore protein ionic current signals, addressing some of the challenges in denoising, segmentation, and amino acid classification. The PELT segmentation produced median segment counts (∼35 per variable region) consistent with the ClpX motor’s reported average 2 amino acid translocation steps, and segment distributions were reproducible across different constructs. We saw a moderate correlation between trace length and segment count, which suggests the algorithm identifies genuine translocation events rather than arbitrarily partitioning signals.

We performed multi-class, multi-group, and pairwise LSTM-based classification of the segmented amino acid region. Individual amino acid region performance varied widely, with negatively charged residues showing consistently strong discrimination across all tasks. In the 20-way classification (with the overall accuracy of ∼21%), D achieved 81.0% accuracy with AUC = 0.97, and E reached 54.1% with AUC = 0.93. This pattern held across binary charge classification (93.3% accuracy, AUC = 0.963), three-way classification (88% recall for negative amino acids), and pairwise comparisons, suggesting that charge state generates highly distinctive ionic current signatures. Pairwise classification (average ∼75% accuracy) showed that many amino acid pairs are readily separable, with confusion concentrated among residues with similar physicochemical properties. Property-based groupings showed that charge produces the strongest discrimination, followed by extreme size differences, while intermediate categories (medium-sized, polar) showed overlapping signal characteristics. We have seen low discrimination among similar amino acids, likely reflecting both biological signal overlap and limited sequence context diversity in the used constructs. The improvement from 20-way to pairwise classification suggests that targeted binary classifiers may be more practical for specific applications like PTM detection or mutation identification rather than attempting simultaneous full amino acid discrimination.

Using dynamic programming segmentation, we segmented traces into fixed 35 segments. These segmented traces were used to make representative templates for each amino acid region using DTW barycenter averaging. We used these profiles for template-trace matching and saw moderate performance. We inspected the DTW alignments of the barycenter against both intra-class and cross-class comparison. We saw good alignments for intra-class comparison. While DTW’s flexible matching properties allow alignment between any two sequences, cross-class comparisons showed either higher costs, increased path deviations, or both compared to intra-class alignments. These suggest that the ionic current signal of different classes can differ enough to cause a higher cost when aligned to traces from other classes. However, the overlap in cost and deviation distributions between intra- and cross-class alignments shows that DTW distance metrics alone provide limited class discrimination for some amino acid pairs.

Using the statistical properties of these barycenters and average variance at each segment, we utilized a profile hidden Markov model for modeling linear segment dependencies and the motor’s stepwise translocation mechanism. This model performed well at coarse-level classification, and its declining discriminative power at finer class resolution may reflect either the constraints of linear-chain modeling or limited sequence context diversity in the current constructs. Profile HMMs are well suited to this setting because they remain trainable on modest amounts of data while yielding interpretable, generative descriptions of translocation, including candidate backslip and stall events. As construct designs diversify and data volume grows, these models can serve as an interpretable baseline and a source of structural priors for deep learning approaches.

## Conclusions

We developed an integrated computational strategy for nanopore protein signal analysis, showing that denoising combined with change point detection algorithms can segment protein translocation events. Segmentation produced consistent results, with segment counts aligning with the ClpX motor’s step size and reproducible performance across different constructs. Classification of the segmented regions showed strong discrimination for charged residues and for property-based groupings, though single-amino-acid resolution across all 20 residues was not achieved with the constructs tested here. This points to practical applications in targeted contexts for differentiating at a coarse-grained level.

Improved performance for challenging amino acid pairs will likely require larger training datasets and optimized construct designs that sample each amino acid across diverse sequence contexts. Future protein constructs incorporating systematic variation in neighboring sequences would enable better characterization of context-dependent signal variation and improve model generalization for amino acids currently showing overlapping signatures. This strategy is also well-suited for the detection of post-translational modifications that alter charge states, identification of specific mutations involving highly discriminable amino acids, or classification tasks focused on distinguishing amino acid groups rather than individual residues.

## Declarations

### Ethics approval and consent to participate

NA

### Consent for publication

All authors have read and approved the manuscript.

## Availability of data and materials

All of the data for this work were achieved from Motone et al.[25]. All the scripts, codes and algorithms used in this study are publicly available in the GitHub repository at https://github.com/genometechlab/nano-protein-signal.

## Author Contributions

**N.G.E:** Data curation, Formal analysis, Methodology, Validation, Visualization, Writing original draft, review & editing. **A.J.S:** Data curation, Formal analysis, Methodology, Validation, Visualization, Writing review & editing. **D.K.-H:** Data curation, Writing review. **J.N.:** Supervision, Writing editing. **M.J:** Conceptualization, Methodology, Project administration, Supervision, Validation, Writing review & editing.

