## Supplementary Information for "A strategy for ionic current analysis of nanopore protein readouts"

### Supplementary Method

Raw Signal processing details: We chose to utilize Huber regression for its ability to be more robust to outliers than median interpolation or any other type of regression, especially considering the points that are ultimately to be averaged into the signal are being identified as outliers. We initialized the isolation forest algorithm with a window size of 15 samples and a contamination parameter of 0.10. This parameter is the prior expectation for how many anomalous points may occur in a signal window. 0.10 or 10% represents that 1-2 points on average may be anomalous in our 15-sample window. The TV denoising algorithm minimizes an objective function that works to: i. Maintain fidelity to the original signal; ii. Optimize against a penalty on the signal's variation. This smooths local fluctuation about the signal mean while maintaining sharp transitions. For this algorithm, we set the regularization parameter (λ) to 0.10. We performed CWT using the Mexican Hat (Ricker) wavelet, whose shape is optimally suited for finding transient peaks in the signal. The transform was computed across logarithmically spaced scales from 1 to 128 samples, providing multi-resolution analysis of spike events. Here, we calculate the spike energy as the sum of the squared coefficients across the high-frequency scales (scales < 5).

**Algorithm 1:** CustomCost-Error(A, start, end)

**Input:** A signal $A$ of length $n$, start index start, end index end

**Output:** The computed cost of the segment $A[\text{start} : \text{end}]$

$\mu \leftarrow \frac{1}{n}\sum_{i=1}^{n} A[i]$ ; `// Compute the mean of the entire signal`

$A \leftarrow A - \mu$ ; `// Center the signal by subtracting the mean`

$S \leftarrow \sum_{i=\text{start}}^{\text{end}} A[i]$ ; `// Compute the sum of the segment`

$Q \leftarrow \sum_{i=\text{start}}^{\text{end}} A[i]^2$ ; `// Compute the sum of squared values in the segment`

$n_{\text{seg}} \leftarrow \text{end} - \text{start} + 1$ ; `// Compute the segment length`

$\text{cost} \leftarrow \text{scale} \times \left(Q - \frac{S^2}{n_{\text{seg}}}\right)$ ; `// Compute the cost based on variance`

**Return** cost ; `// Return the computed cost for the segment`

### Supplementary Figures

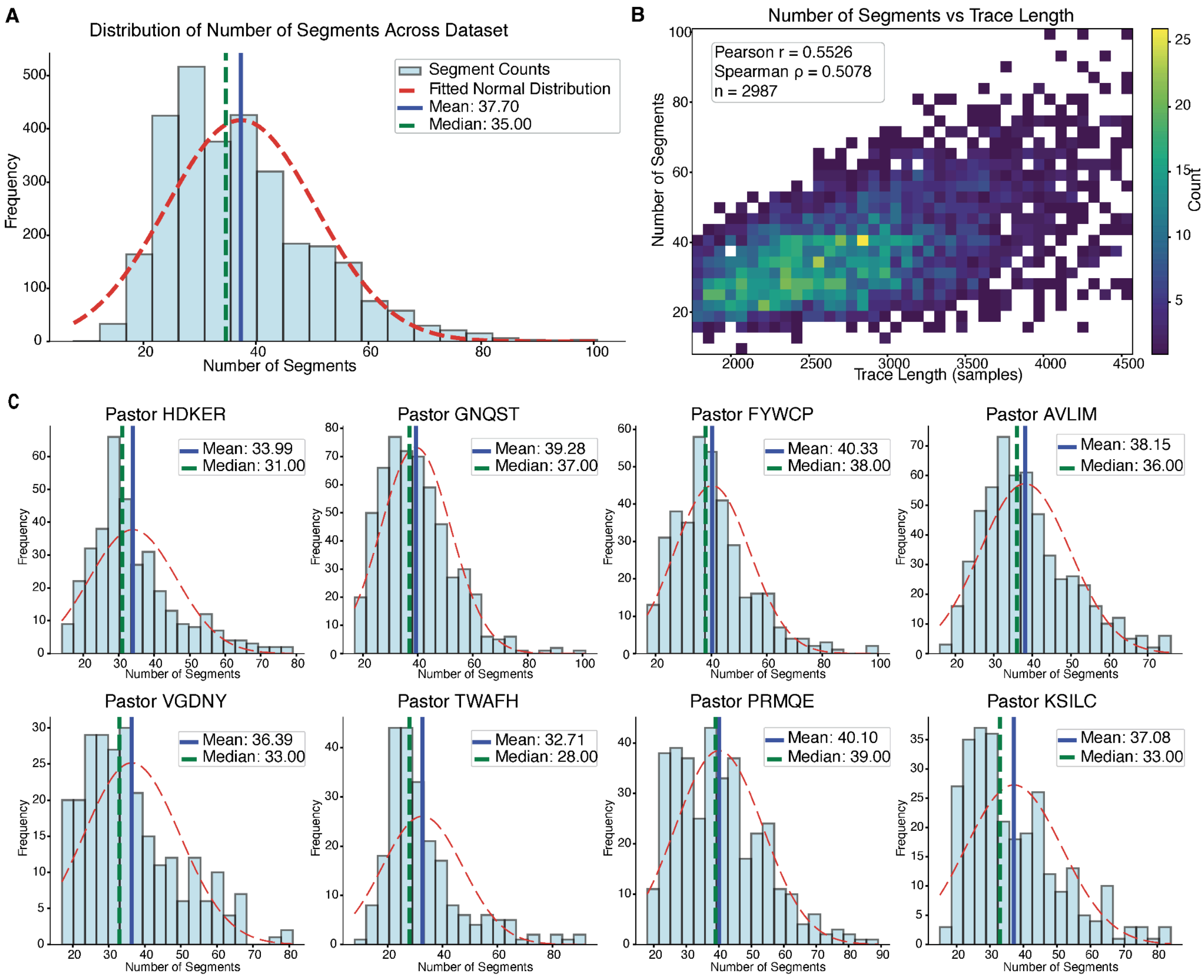


**Supplementary Figure 1. Distribution of segment counts from PELT segmentation across all variable regions**. **A.** Overall distribution of segment counts across the entire dataset (n=2,987 variable regions). Histogram shows frequency of traces with different numbers of segments. Red dashed curve represents fitted normal distribution. Blue line indicates the mean (37.70 segments), and green dashed line shows the median (35.00 segments). **B.** Relationship between trace length and number of segments. Heatmap showing the joint distribution of trace length (x-axis, in samples) and resulting segment count (y-axis). Color intensity indicates the number of traces in each bin, with warmer colors representing higher density. Pearson correlation coefficient r = 0.5526 and Spearman rank correlation ρ = 0.5078 indicate moderate positive correlation between trace length and segment count (n=2,987). **C.** Segment count distributions for individual PASTOR constructs. Histograms show the distribution of segment counts for each of the eight PASTOR variants: HDKER, GNQST, FYWCP, AVLIM, VGDNY, TWAFH, PRMQE, and KSILC. Blue lines indicate mean segment counts, green dashed lines show median values, and red dashed curves represent fitted normal distributions for each PASTOR. All constructs show similar distributional characteristics with means ranging from 32.71 to 40.33 segments and medians from 28.00 to 39.00 segments.

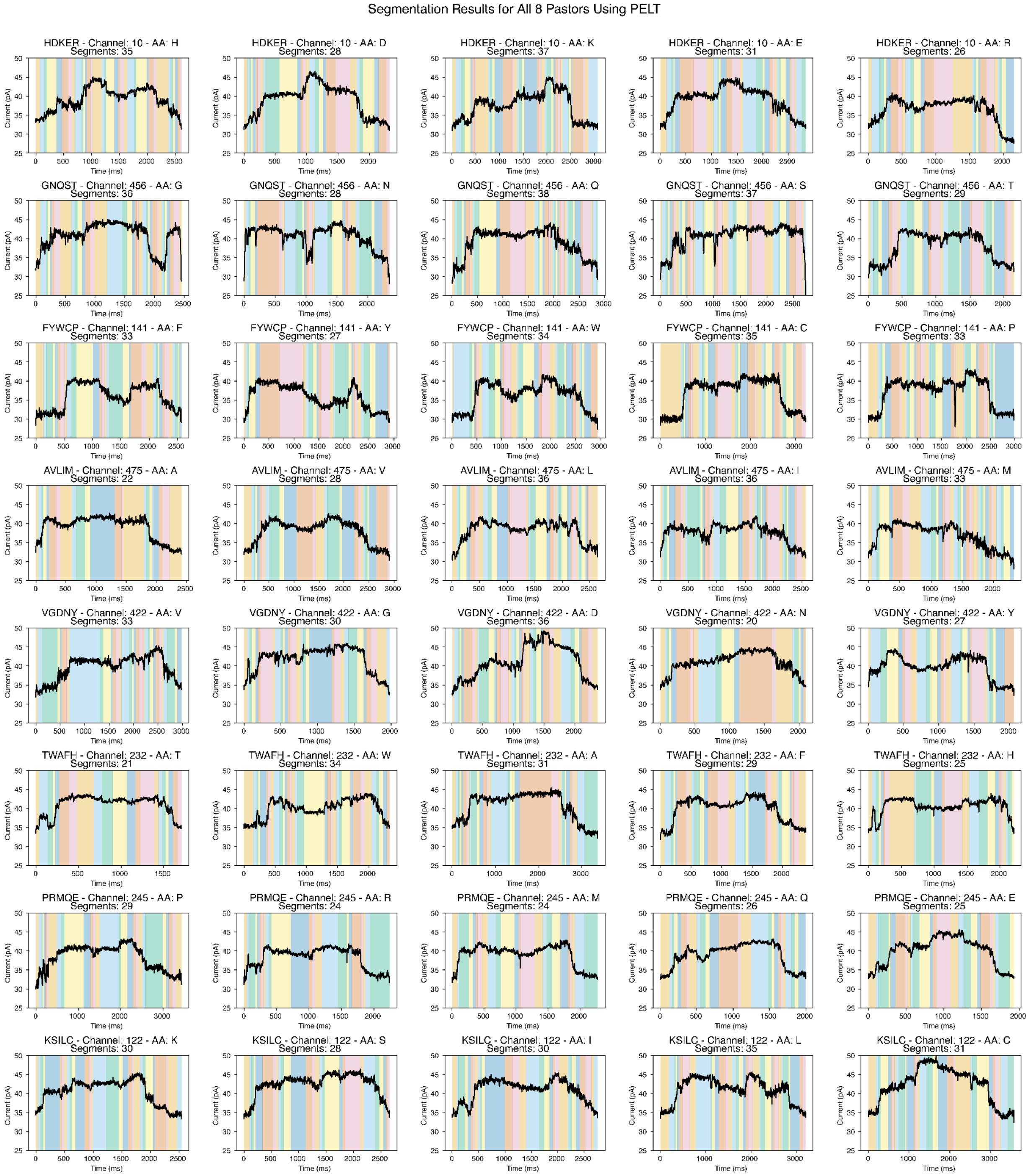


**Supplementary Figure 2. PELT segmentation across all 8 PASTOR constructs.** Representative PELT segmentation results for all five amino acid variable regions from each of the eight PASTOR constructs (HDKER, GNQST, FYWCP, AVLIM, VGDNY, TWAFH, PRMQE, KSILC). Each panel shows a single variable region trace (black line) with colored vertical bands indicating detected segments. Titles include PASTOR name, channel number, amino acid label, and total segment count. All traces were segmented using penalty=5, min_size=30, and custom variance-based cost function.

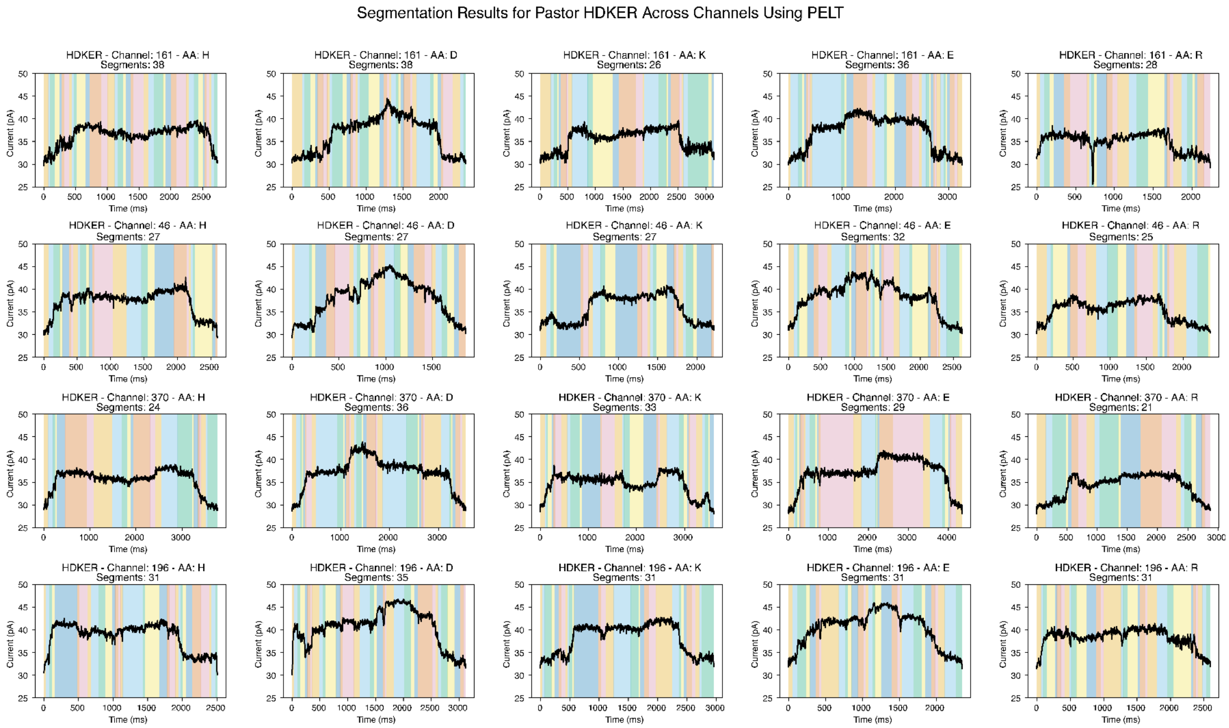


**Supplementary Figure 3. PELT segmentation across the same constructs.** Representative PELT segmentation results for all five amino acid variable regions from three examples of the same construct (HDKER). Each panel shows a single variable region trace (black line) with colored vertical bands indicating detected segments. Titles include PASTOR name, channel number, amino acid label, and total segment count. All traces were segmented using penalty=5, min_size=30, and custom variance-based cost function.

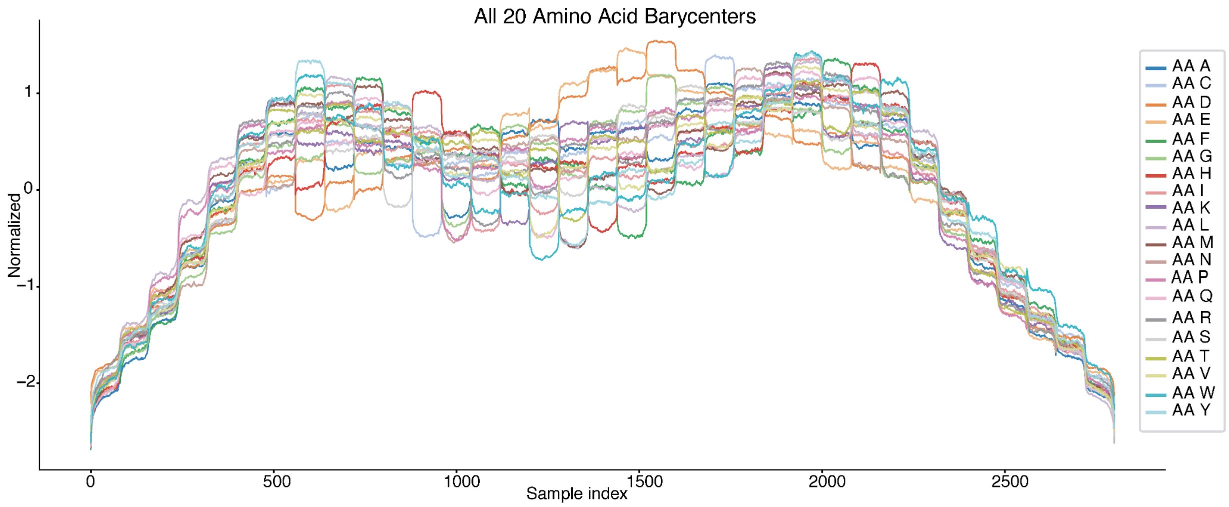


**Supplementary Figure 4. DTW barycenters for all 20 amino acids.** All 20 amino acid barycenters overlaid, each constructed from Dynamic Programming-segmented traces (35 segments per trace) using DTW barycenter averaging. Each barycenter represents the consensus ionic current pattern for one amino acid class, computed from all available traces after z-score normalization and interpolation to 80 points per segment. The x-axis shows concatenated sample index across all 35 segments (2,800 total points), and the y-axis shows normalized current.

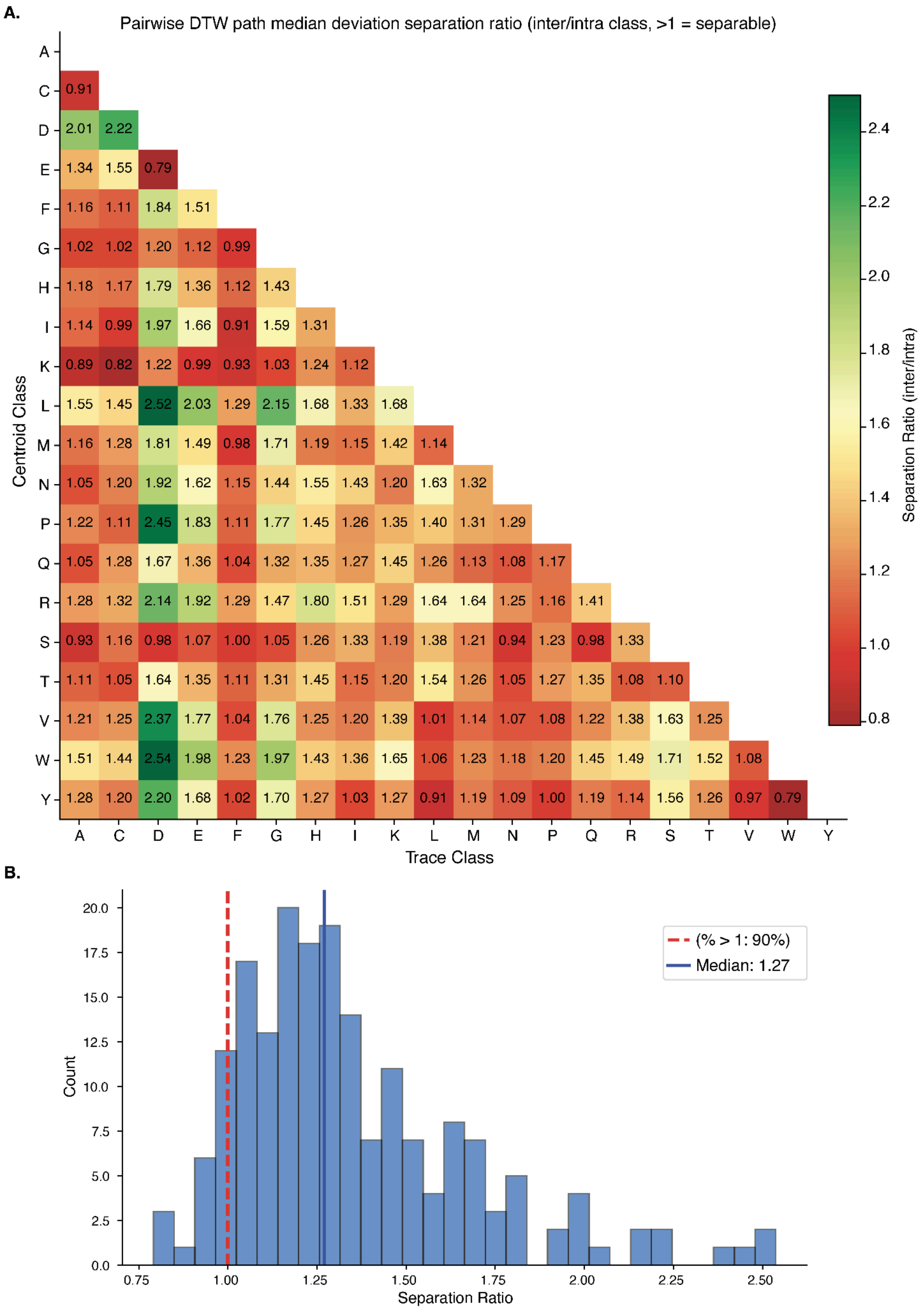


**Supplementary Figure 5.Pairwise barycenter separation analysis using DTW path deviation. A.** Separation ratio matrix for all 190 amino acid pairs. Each cell (i,j) shows the ratio of median inter-class distance (centroid i to traces j) divided by median intra-class distance (centroid i to traces i). Only the lower triangle is shown to avoid redundancy. Values indicate exact separation ratios. **B.** Distribution of separation ratios across all 190 pairs. Red dashed line at ratio=1.0 indicates the threshold for separation. Approximately 90% of pairs showed ratios >1.0.

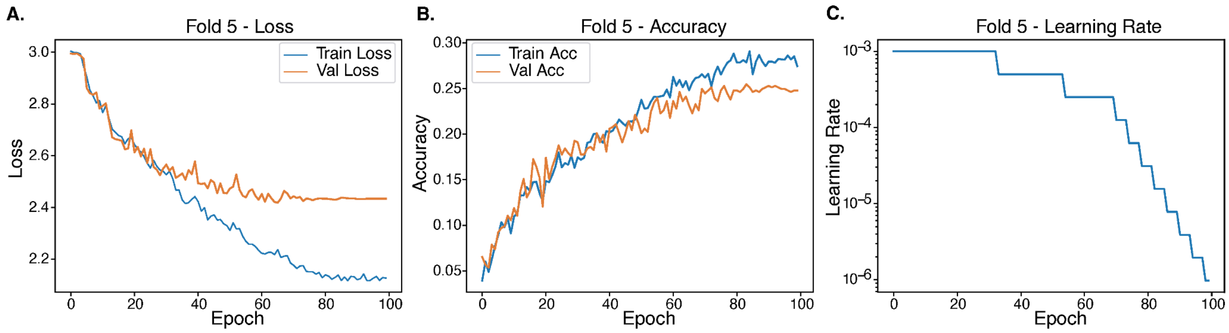


**Supplementary Figure 6. Training dynamics for LSTM 20-way amino acid classification (Fold 5)**. **A.** Training and validation loss curves over 100 epochs. Both losses decrease steadily, with validation loss plateauing around epoch 70 at approximately 2.4. **B.** Training and validation accuracy curves over 100 epochs. Both accuracies increase from ~5% to stabilize around 25-29% in later epochs. Close tracking between training and validation curves shows learning of generalizable features. **C.** Learning rate schedule over training. The adaptive learning rate scheduler progressively reduced the learning rate from $10^{-3}$ to ~$10^{-6}$, with reductions triggered when validation loss plateaued, enabling fine-tuned optimization in later training phases.

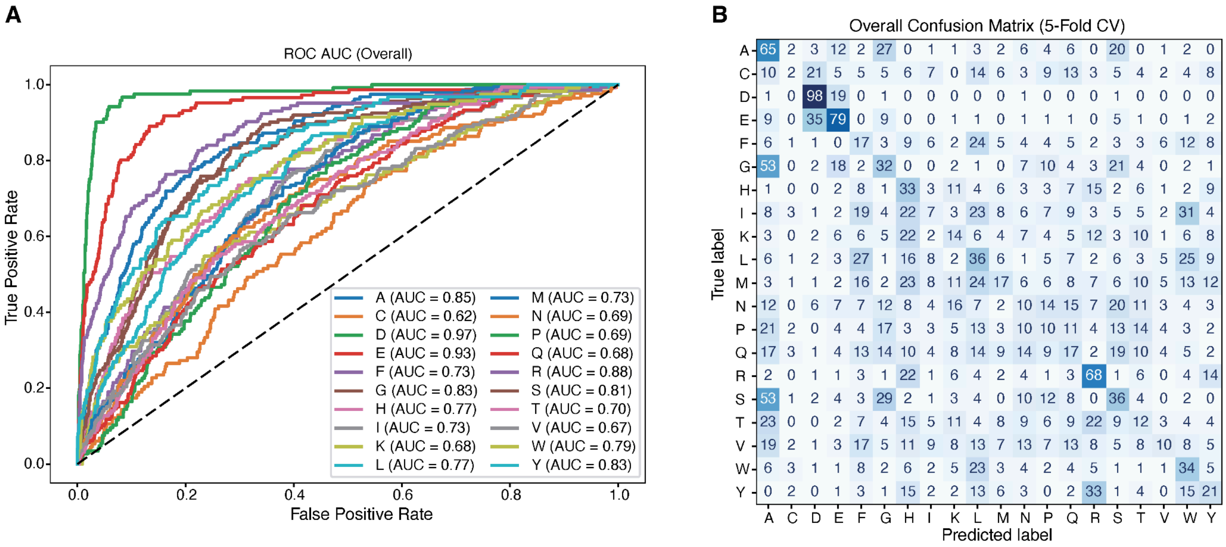


**Supplementary Figure 7. Performance metrics for 20-way amino acid classification using bidirectional LSTM. A.** One-vs-rest ROC curves for all 20 amino acid classes. Each curve represents the classifier's ability to distinguish one amino acid from all others. AUC values range from 0.62 (C) to 0.97 (D). Dashed line indicates random classifier performance (AUC = 0.5). **B.** Confusion matrix showing predicted versus true labels across 5-fold cross-validation (n=2,987 total predictions). Rows represent true amino acid labels, columns represent predicted labels. Diagonal cells show correct classifications, while off-diagonal cells indicate misclassifications. Numbers in each cell indicate prediction counts.

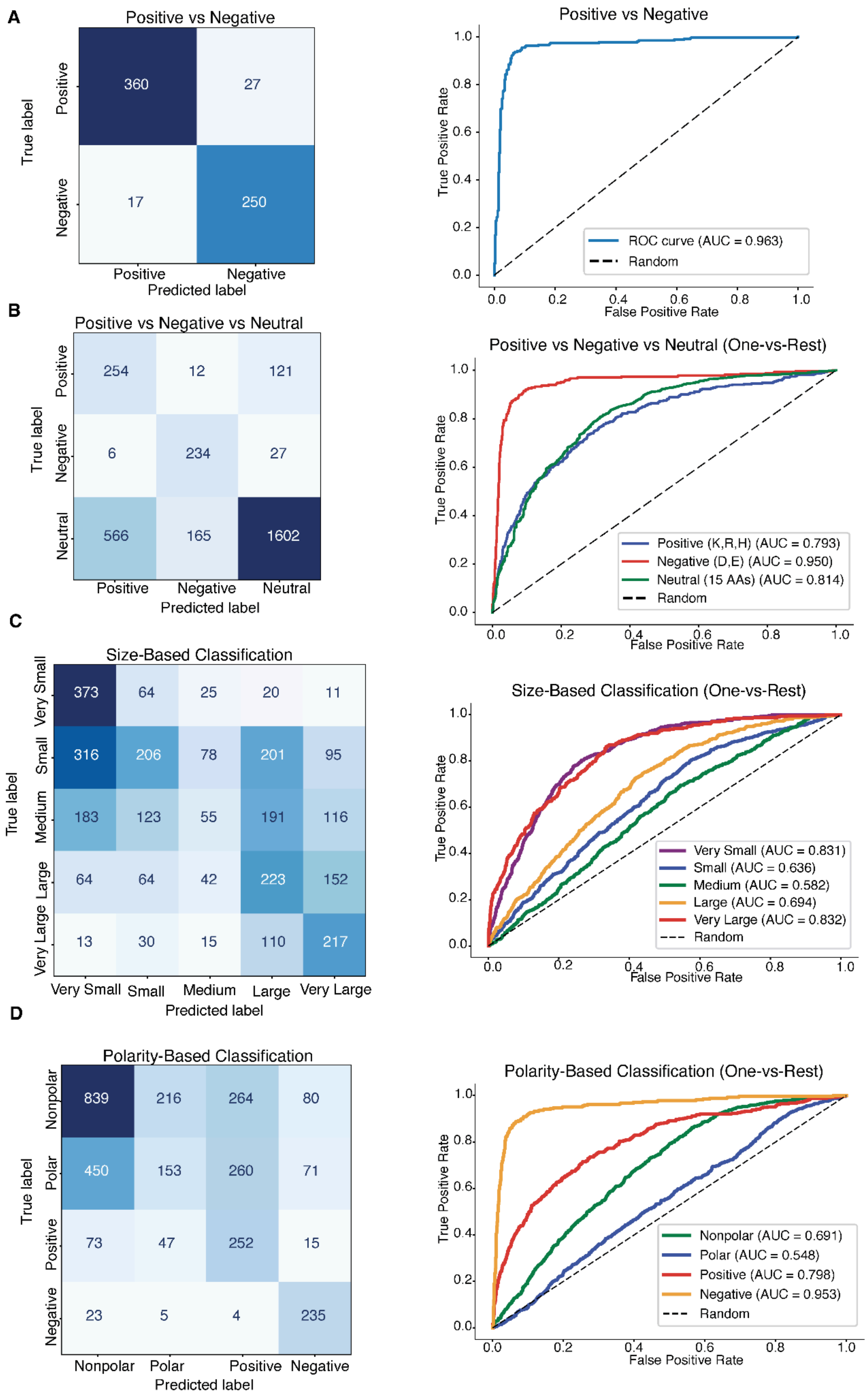
A
Positive vs Negative
True label
Positive
Negative
360
27
17
250
Predicted label
Positive vs Negative
True Positive Rate
False Positive Rate
ROC curve (AUC = 0.963)
Random
B
Positive vs Negative vs Neutral
True label
Positive
Negative
Neutral
254
12
121
6
234
27
566
165
1602
Predicted label
Positive vs Negative vs Neutral (One-vs-Rest)
True Positive Rate
False Positive Rate
Positive (K,R,H) (AUC = 0.793)
Negative (D,E) (AUC = 0.950)
Neutral (15 AAs) (AUC = 0.814)
Random
C
Size-Based Classification
True label
Very Small
Small
Medium
Large
Very Large
373
64
25
20
11
316
206
78
201
95
183
123
55
191
116
64
64
42
223
152
13
30
15
110
217
Predicted label
Size-Based Classification (One-vs-Rest)
True Positive Rate
False Positive Rate
Very Small (AUC = 0.831)
Small (AUC = 0.636)
Medium (AUC = 0.582)
Large (AUC = 0.694)
Very Large (AUC = 0.832)
Random
D
Polarity-Based Classification
True label
Nonpolar
Polar
Positive
Negative
839
216
264
80
450
153
260
71
73
47
252
15
23
5
4
235
Predicted label
Polarity-Based Classification (One-vs-Rest)
True Positive Rate
False Positive Rate
Nonpolar (AUC = 0.691)
Polar (AUC = 0.548)
Positive (AUC = 0.798)
Negative (AUC = 0.953)
Random

**Supplementary Figure 8. Multi-group LSTM classification performance based on physicochemical properties. A.** Binary charge-based classification (Positive vs. Negative). Left: Confusion matrix showing classification of positively charged amino acids (K, R, H; n=387) versus negatively charged amino acids (D, E; n=267). Numbers indicate the count of samples and predictions in each category, with darker colors indicating higher counts. Right: ROC curve for binary classification with AUC = 0.963. Dashed line represents random classifier performance. **B.** Three-way charge-based classification (Positive vs. Negative vs. Neutral). Left: Confusion matrix for classification of positively charged (K, R, H; n=387), negatively charged (D, E; n=267), and neutral amino acids (15 remaining amino acids; n=2,333). Right: One-vs-rest ROC curves for each class. Positive amino acids shown in blue (AUC = 0.793), negative amino acids in red (AUC = 0.950), and neutral amino acids in green (AUC = 0.814). Dashed line represents random classifier. **C.** Five-way size-based classification. Left: Confusion matrix for classification based on molecular size: Very Small (G, A, S; n=493), Small (C, D, N, P, T, V; n=896), Medium (E, I, L, Q; n=668), Large (H, K, M, F; n=545), and Very Large (R, W, Y; n=385). Right: One-vs-rest ROC curves for each size category. Very Small in purple (AUC = 0.831), Small in blue (AUC = 0.636), Medium in green (AUC = 0.582), Large in orange (AUC = 0.694), and Very Large in red (AUC = 0.832). Dashed line represents random classifier. **D.** Four-way polarity-based classification. Left: Confusion matrix for classification by polarity: Nonpolar (A, G, I, L, M, F, W, P, V; n=1,399), Polar (S, T, N, Q, C, Y; n=934), Positive (K, R, H; n=387), and Negative (D, E; n=267). Right: One-vs-rest ROC curves for each polarity category. Nonpolar in green (AUC = 0.691), Polar in blue (AUC = 0.548), Positive in green (AUC = 0.798), and Negative in orange (AUC = 0.953). Dashed line represents a random classifier.

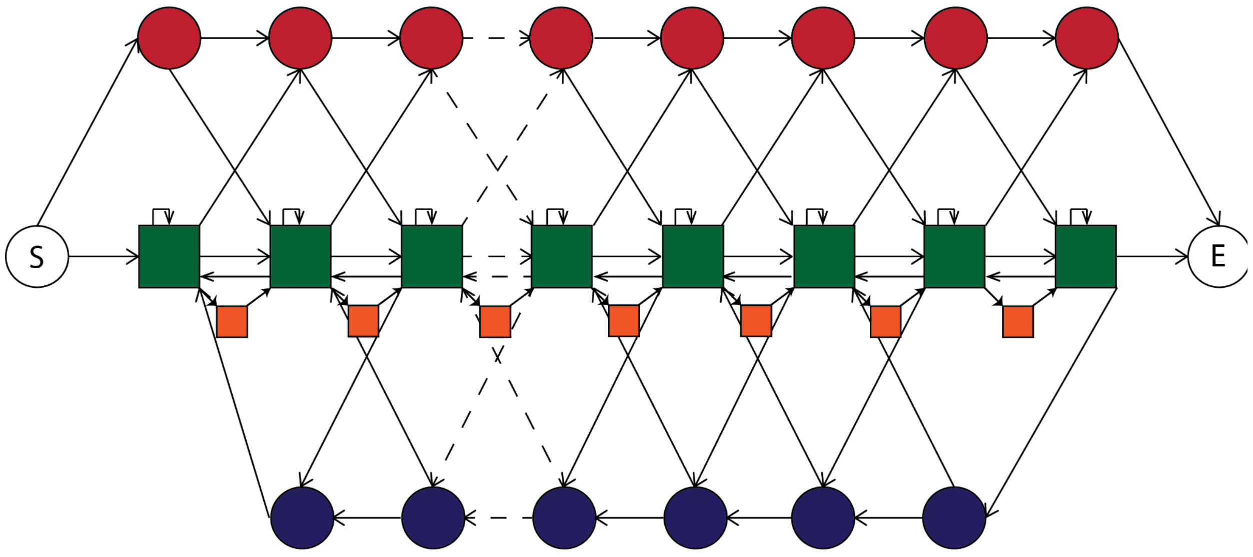


**Supplementary Figure 9. Structure of profile hidden Markov model.** Each square state represents an emission, while circular states represent silent model traversal states. The green squares are the input profiles for every amino acid, assigned with their barycenter means and segment-variance pileup. The orange squares are the average states between each flanking profile state. The red circle represents a silent deletion state which can skip over profile states at an exponential cost. The blue circle represents an insert state that can continuously insert anywhere at an exponential cost. Both S and E represent silent start and end states the model needs to continue. Every arrow represents a possible transition that can be made by the model.

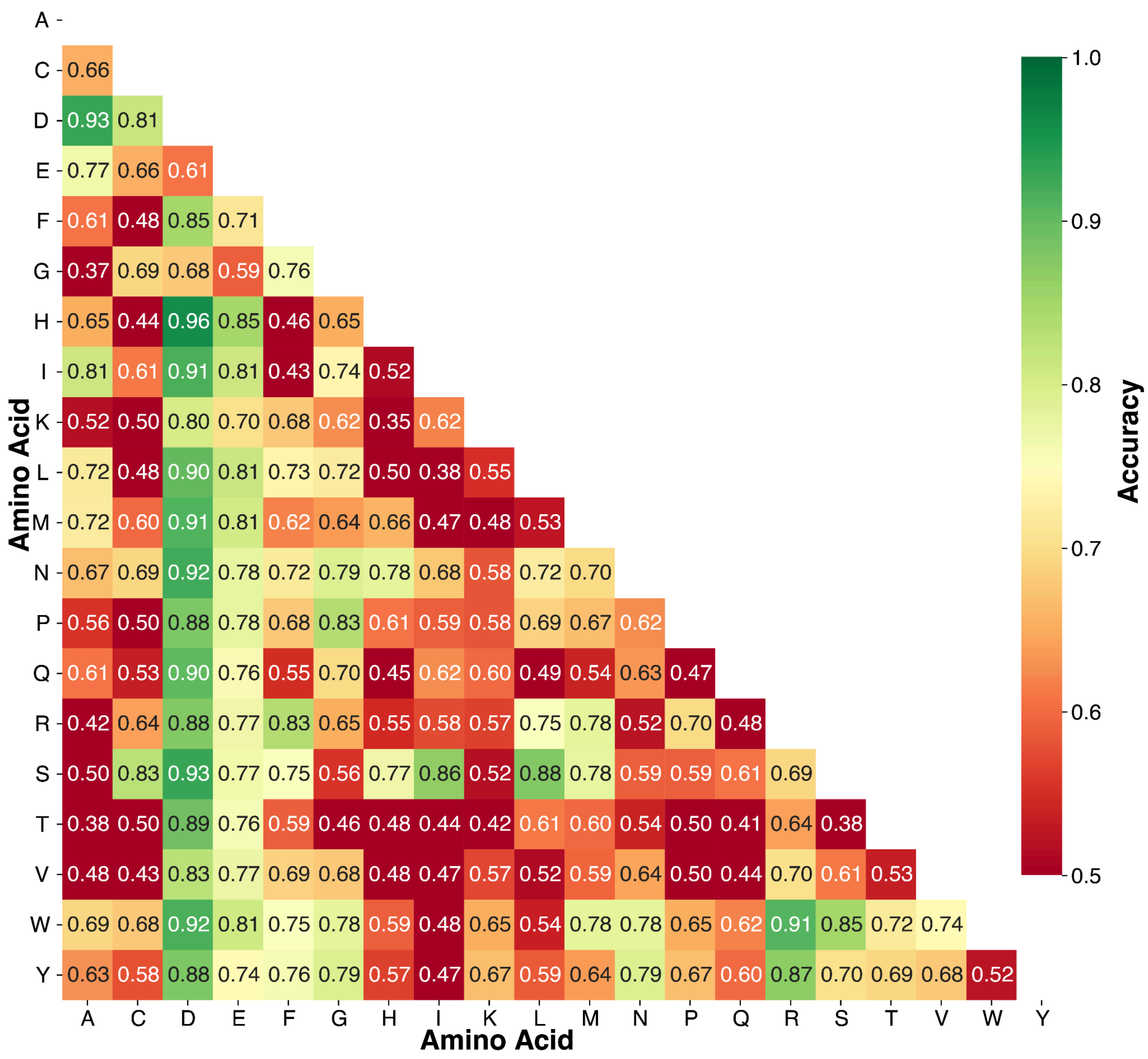


**Supplementary Figure 10. Pairwise HMM Classification Matrix.** A subset of traces were split into training and test sets. The training set was used with dynamic barycenter averaging to create a profile, where the mean is calculated from the DBA profile, and the variance is calculated from the first 10 segments in the test set that align at that position. These features are used in the profile hidden Markov model in order to get a log-probability for every trace when compared to a given amino acid profile. These pairwise classifications are shown within this plot, where the accuracy of classifying an amino acid from another amino acid is shown.

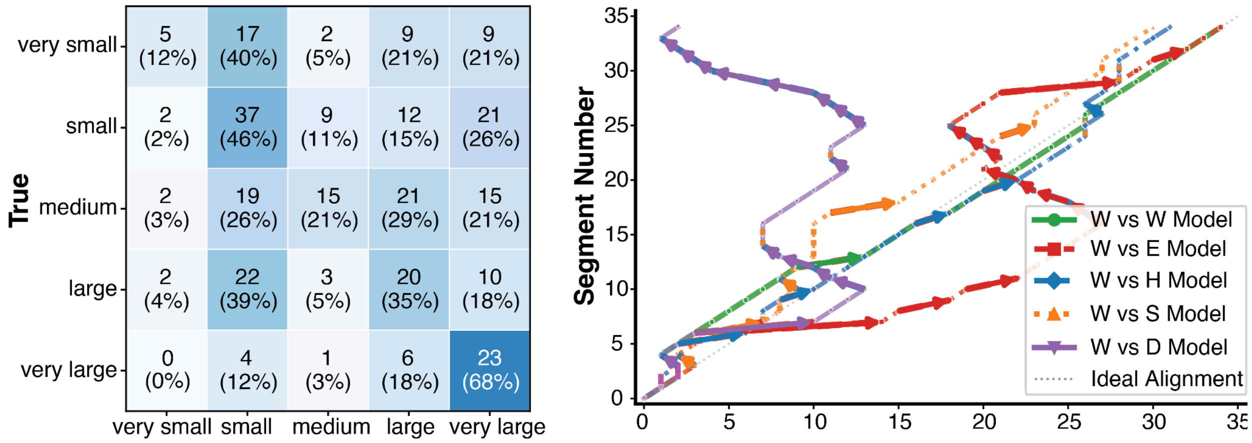


**Supplementary Figure 11. Hidden Markov model inference. Left:** Five-way size-based classification. Confusion matrix for classification based on molecular size: Very Small (G, A, S; n=42), Small (C, D, N, P, T, V; n=81), Medium (E, I, L, Q; n=72), Large (H, K, M, F; n=57), and Very Large (R, W, Y; n=34). **Right:** Model traversal of amino acid W vs. its own model (green), and W vs. E (red), W vs. H (blue), W vs. S (orange), and W vs. D (purple).

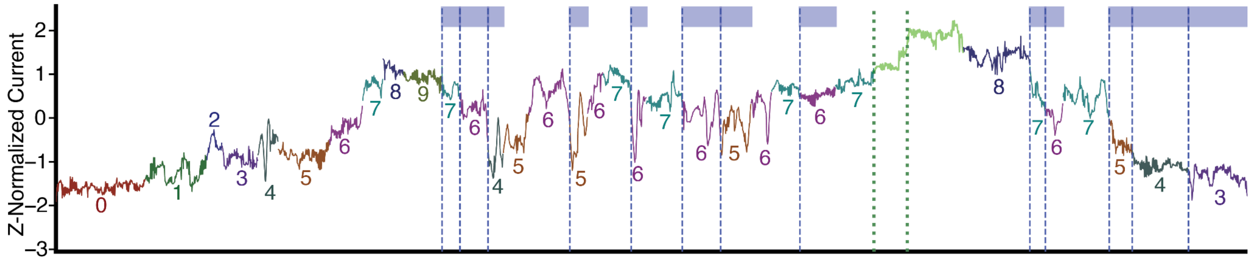


**Supplementary Figure 12. HMM Inference on Segmented Ionic Current Traces.** Segmented ionic current for Proline (P). The count and color sequence represents each unique match state the model traversed when inferring the trace. The red dashed line represents a point where the model chose to skip model match states until a downstream match state. The blue dashed line represents a point where the model slipped backwards to an upstream match state. The green dashed line and following segment represents the model inserting a state with a mean and variance averaged between the preceding and following states. The y-axis is z-score normalized ionic current in picoamperes.

### Supplementary Tables

**Table S1.** Implementation details and parameters for the ionic current signal processing framework.

| **Parameter** | **Value** |
|---|---|
| **Signal preprocessing and denoising** | |
| Anomaly detection algorithm | Isolation Forest |
| Isolation Forest, number of estimators | 100 |
| Isolation Forest, contamination threshold | 0.10 |
| Noise reduction algorithm | Total variation denoising |
| Total variation denoising, regularization weight | 0.10 |
| Spike detection algorithm | Continuous wavelet transform |
| Continuous wavelet transform, wavelet family and scales | Mexican Hat (Ricker) waves, logarithmically spaces scales from 1 to 128. |
| Spike threshold | median + 0.25 × MAD |
| Spike dilation | ±1 sample |
| Signal reconstruction | Huber regression |
| Huber regression, epsilon and alpha | 2.0; 0.01 |
| Variable region boundary assignment | absolute minimum of each tyrosine-tyrosine dip |
| **Segmentation** | |
| Search methods evaluated | binary segmentation, window-based, bottom-up, PELT, Dynp |
| Search methods retained | PELT (variable number), Dynp (fixed number) |
| Cost function | custom variance-based (Algorithm S1) |
| PELT, penalty (β) | 5 |
| PELT, minimum segment size | 30 samples |
| Dynp, number of breakpoints | 34 (35 segments per trace) |
| Dynp, minimum segment size | 30 |
| Implementation | ruptures |
| **Dynamic time warping and barycenter averaging** | |
| Trace normalization | z-score, per trace |
| Segment interpolation | linear, to 80 samples |
| Template dimensionality | 35 segments × 80 samples |
| Barycenter algorithm | DTW barycenter averaging (DBA) |

| | |
|---|---|
| DTW distance normalization | path cost ÷ mean of the two segment lengths |
| Alignment metrics | total path cost; mean cost per aligned segment pair; mean $\|i - j\|$ deviation from diagonal |
| Separation ratio | median inter-class deviation ÷ median intra-class deviation |
| Implementation | tslearn, |
| **Bidirectional LSTM classifier** | |
| Segment features | mean, median, standard deviation, minimum, maximum, skewness, kurtosis, segment length |
| Architecture | two-layer bidirectional LSTM |
| Hidden units | 64 per direction |
| Variable-length sequence handling | packed sequences |
| Recurrent dropout | 0.30 |
| Dense head | 128 units, ReLU activation, dropout 0.30 |
| Output units | 20 (multiclass); 2 (pairwise and binary charge); 3, 4 or 5 (group schemes) |
| Loss | cross-entropy, class weights inverse to class frequency |
| Optimizer | AdamW |
| Learning rate | 0.001 |
| Weight decay | 0.0001 |
| Learning rate schedule | ×0.5 on validation loss plateau, patience 3 |
| Gradient clipping, maximum norm | 1.0 |
| Batch size | 32, padded within batch |
| Epochs | 100 |
| Multiclass and group evaluation | stratified five-fold cross-validation, fold predictions pooled |
| Pairwise evaluation | 190 binary classifiers, 80/20 stratified split |
| Implementation | PyTorch |
| **Profile hidden Markov model** | |
| Package | YAHMM |
| Topology | linear profile of emitting and silent states |
| Match state emission | Gaussian |
| Match state mean | DBA segment mean at that position |

| | |
|---|---|
| Match state variance | pileup variance across traces at that position |
| Traces contributing to variance estimate | 10 |
| Skip state | silent; forward transition to a downstream match state (deletion) |
| Slip state | silent; backward transition to an upstream match state (insertion or repeat) |
| Interpolated state | emission parameters averaged from the flanking profile states |
| Additional transitions | self-loops on match states |
| Trace filter | interquartile range of trace length |
| Data split | 60% profile construction, 40% inference |
| Inference | Viterbi decoding; segment means scored as univariate log-odds |
| Model assignment | highest-scoring model |
| Label schemes evaluated | pairwise; charge (two- and three-way); size (five-way); polarity (four-way) |

| **Table S2. LSTM 20-Way Amino Acid Classification** | | | | | | |
|---|---|---|---|---|---|---|
| Class | Per-Class Accuracy | Precision | Recall | F1-Score | ROC AUC | Support |
| A (0) | 0.414 | 0.20 | 0.41 | 0.27 | 0.85 | 157 |
| C (1) | 0.015 | 0.09 | 0.02 | 0.03 | 0.62 | 132 |
| D (2) | 0.810 | 0.55 | 0.81 | 0.66 | 0.97 | 121 |
| E (3) | 0.541 | 0.45 | 0.54 | 0.49 | 0.93 | 146 |
| F (4) | 0.141 | 0.10 | 0.14 | 0.12 | 0.73 | 121 |
| G (5) | 0.198 | 0.18 | 0.20 | 0.19 | 0.83 | 162 |
| H (6) | 0.282 | 0.15 | 0.28 | 0.19 | 0.77 | 117 |
| I (7) | 0.041 | 0.10 | 0.04 | 0.06 | 0.73 | 172 |
| K (8) | 0.111 | 0.13 | 0.11 | 0.12 | 0.68 | 126 |
| L (9) | 0.211 | 0.16 | 0.21 | 0.18 | 0.77 | 171 |
| M (10) | 0.094 | 0.18 | 0.09 | 0.12 | 0.73 | 181 |

| | | | | | | |
|---|---|---|---|---|---|---|
| N (11) | 0.060 | 0.08 | 0.06 | 0.07 | 0.69 | 168 |
| P (12) | 0.069 | 0.09 | 0.07 | 0.08 | 0.69 | 146 |
| Q (13) | 0.095 | 0.12 | 0.09 | 0.10 | 0.68 | 179 |
| R (14) | 0.472 | 0.35 | 0.47 | 0.40 | 0.88 | 144 |
| S (15) | 0.207 | 0.20 | 0.21 | 0.20 | 0.81 | 174 |
| T (16) | 0.076 | 0.10 | 0.08 | 0.09 | 0.70 | 157 |
| V (17) | 0.058 | 0.21 | 0.06 | 0.09 | 0.67 | 172 |
| W (18) | 0.291 | 0.19 | 0.29 | 0.23 | 0.79 | 117 |
| Y (19) | 0.169 | 0.18 | 0.17 | 0.17 | 0.83 | 124 |
| Macro Average | - | 0.19 | 0.22 | 0.19 | 0.76 | 2987 |
| Weighted Average | - | 0.19 | 0.21 | 0.19 | 0.77 | 2987 |
| Overall Accuracy | 0.207 ± 0.029 | - | - | - | - | 2987 |

**Table S3. LSTM Binary Charge-Based Classification: Positive vs. Negative**

| Class | Per-Class Accuracy | Precision | Recall | F1-Score | ROC AUC | Support |
|---|---|---|---|---|---|---|
| Positive (K,R,H) | 0.930 | 0.95 | 0.93 | 0.94 | - | 387 |
| Negative (D,E) | 0.936 | 0.90 | 0.94 | 0.92 | - | 267 |
| Macro Average | - | 0.93 | 0.93 | 0.93 | - | 654 |
| Weighted Average | - | 0.93 | 0.93 | 0.93 | - | 654 |
| Overall Accuracy | 0.933 ± 0.013 | - | - | - | 0.963 | 654 |

**Table S4. LSTM Three-Way Charge-Based Classification: Positive vs. Negative vs. Neutral**

| Class | Per-Class Accuracy | Precision | Recall | F1-Score | ROC AUC | Support |
|---|---|---|---|---|---|---|
| Positive (K,R,H) | 0.656 | 0.31 | 0.66 | 0.42 | 0.793 | 387 |

| | | | | | | |
|---|---|---|---|---|---|---|
| Negative (D,E) | 0.876 | 0.57 | 0.88 | 0.69 | 0.950 | 267 |
| Neutral (15 AAs) | 0.687 | 0.92 | 0.69 | 0.78 | 0.814 | 2333 |
| Macro Average | - | 0.60 | 0.74 | 0.63 | 0.852 | 2987 |
| Weighted Average | - | 0.81 | 0.70 | 0.73 | 0.823 | 2987 |
| Overall Accuracy | 0.700 ± 0.019 | - | - | - | - | 2987 |

**Table S5. LSTM Five-Way Size-Based Classification**

| Class | Per-Class Accuracy | Precision | Recall | F1-Score | ROC AUC | Support |
|---|---|---|---|---|---|---|
| Very Small (G,A,S) | 0.757 | 0.39 | 0.76 | 0.52 | 0.831 | 493 |
| Small (C,D,N,P,T,V) | 0.230 | 0.42 | 0.23 | 0.30 | 0.636 | 896 |
| Medium (E,I,L,Q) | 0.082 | 0.26 | 0.08 | 0.12 | 0.582 | 668 |
| Large (H,K,M,F) | 0.409 | 0.30 | 0.41 | 0.35 | 0.694 | 545 |
| Very Large (R,W,Y) | 0.564 | 0.37 | 0.56 | 0.44 | 0.832 | 385 |
| Macro Average | - | 0.35 | 0.41 | 0.35 | 0.715 | 2987 |
| Weighted Average | - | 0.35 | 0.36 | 0.32 | 0.692 | 2987 |
| Overall Accuracy | 0.360 ± 0.016 | - | - | - | - | 2987 |

**Table S6. LSTM Four-Way Polarity-Based Classification**

| Class | Per-Class Accuracy | Precision | Recall | F1-Score | ROC AUC | Support |
|---|---|---|---|---|---|---|
| Nonpolar (A,G,I,L,M,F,W,P,V) | 0.600 | 0.61 | 0.60 | 0.60 | 0.691 | 1399 |
| Polar (S,T,N,Q,C,Y) | 0.164 | 0.36 | 0.16 | 0.23 | 0.548 | 934 |
| Positive (K,R,H) | 0.651 | 0.32 | 0.65 | 0.43 | 0.798 | 387 |
| Negative (D,E) | 0.880 | 0.59 | 0.88 | 0.70 | 0.953 | 267 |
| Macro Average | - | 0.47 | 0.57 | 0.49 | 0.747 | 2987 |

| | | | | | | |
|---|---|---|---|---|---|---|
| Weighted Average | - | 0.49 | 0.50 | 0.47 | 0.683 | 2987 |
| Overall Accuracy | 0.495 ± 0.015 | - | - | - | - | 2987 |

**Table 6. HMM State Traceback for two-way classification**

| Category | Traces | Match Mean | Match Std | Backslip Mean | Backslip Std | Skip Mean | Skip Std | Self Loop Mean | Self Loop Std | Insert Mean | Insert Std |
|---|---|---|---|---|---|---|---|---|---|---|---|
| Positive | 34 | 28.82 | 5.58 | 2.29 | 3.44 | 6.24 | 2.14 | 0.15 | 0.43 | 1.65 | 1.33 |
| Negative | 28 | 27.71 | 3.95 | 2.82 | 2.67 | 8.54 | 2.99 | 0.07 | 0.26 | 2.11 | 1.42 |

**Table 7. HMM State Traceback for three-way classification**

| Class | Traces | Match Mean | Match Std | Backslip Mean | Backslip Std | Skip Mean | Skip Std | Self Loop Mean | Self Loop Std | Insert Mean | Insert Std |
|---|---|---|---|---|---|---|---|---|---|---|---|
| Positive | 34 | 28.82 | 5.58 | 2.29 | 3.44 | 6.24 | 2.14 | 0.15 | 0.43 | 1.65 | 1.33 |
| Negative | 28 | 27.71 | 3.95 | 2.82 | 2.67 | 8.54 | 2.99 | 0.07 | 0.26 | 2.11 | 1.42 |
| Neutral | 224 | 29.25 | 4.57 | 2.03 | 2.78 | 6.33 | 3.21 | 0.11 | 0.35 | 1.75 | 1.29 |

**Table 8. HMM State Traceback for four-way polarity-based classification**

| Class | Traces | Match Mean | Match Std | Backslip Mean | Backslip Std | Skip Mean | Skip Std | Self Loop Mean | Self Loop Std | Insert Mean | Insert Std |
|---|---|---|---|---|---|---|---|---|---|---|---|
| Nonpolar | 139 | 28.90 | 5.24 | 2.24 | 3.23 | 6.34 | 3.27 | 0.09 | 0.28 | 1.78 | 1.39 |
| Polar | 85 | 29.84 | 3.11 | 1.67 | 1.76 | 6.32 | 3.10 | 0.14 | 0.44 | 1.71 | 1.10 |
| Positive | 34 | 28.82 | 5.58 | 2.29 | 3.44 | 6.24 | 2.14 | 0.15 | 0.43 | 1.65 | 1.33 |
| Negative | 28 | 27.71 | 3.95 | 2.82 | 2.67 | 8.54 | 2.99 | 0.07 | 0.26 | 2.11 | 1.42 |

**Table 9. HMM State Traceback for five-way size-based classification**

| Class | Traces | Match Mean | Match Std | Backslip Mean | Backslip Std | Skip Mean | Skip Std | Self Loop Mean | Self Loop Std | Insert Mean | Insert Std |
|---|---|---|---|---|---|---|---|---|---|---|---|
| Very Small | 42 | 28.40 | 5.76 | 2.55 | 3.33 | 6.12 | 3.67 | 0.10 | 0.29 | 1.57 | 1.28 |

| Table 9. HMM State Traceback for five-way size-based classification | | | | | | | | | | |
|---|---|---|---|---|---|---|---|---|---|---|
| Small | 81 | 29.48 | 4.47 | 2.02 | 2.83 | 6.17 | 3.12 | 0.15 | 0.45 | 1.58 | 1.13 |
| Medium | 72 | 29.14 | 3.38 | 2.14 | 2.21 | 7.38 | 2.94 | 0.06 | 0.23 | 1.83 | 1.29 |
| Large | 57 | 28.28 | 5.37 | 2.28 | 3.15 | 6.39 | 2.87 | 0.14 | 0.35 | 2.07 | 1.51 |
| Very Large | 34 | 29.94 | 4.43 | 1.65 | 3.00 | 6.38 | 3.07 | 0.09 | 0.37 | 1.88 | 1.32 |
